# Dual Th1 and tissue-repair Treg cells accumulate in skeletal muscle preserving tissue integrity during infection

**DOI:** 10.64898/2026.06.29.734788

**Authors:** Cintia L. Araujo Furlan, Santiago Boccardo, Camila M.S. Gimenez, Yamila Gazzoni, Julio Gareca, Constanza Rodriguez, Jorge H. Mukdsi, María C. Amezcua Vesely, Adriana Gruppi, Carolina L. Montes, Bola S. Hanna, Eva V. Acosta Rodríguez

## Abstract

Chronic infections require mechanisms that limit tissue damage while preserving pathogen control, yet the contribution of regulatory T (Treg) cells to this balance remains unclear. In this study, we characterized Treg cell responses during a chronic parasitic infection using experimental *Trypanosoma cruzi* infection as a model of persistent low-level parasitism and chronic tissue inflammation. We found that, although Treg cell numbers decline in the spleen, they accumulate in parasite-affected tissues such as skeletal muscle, where they adopt a combined Th1-associated and tissue-repair program. Systemic Treg cell depletion had limited impact on immune and disease-associated parameters, whereas local depletion in skeletal muscle exacerbated tissue damage and increased parasite burden. Moreover, transient systemic perturbation of Treg cells during the acute phase impaired their long-term accumulation in skeletal muscle, resulting in increased tissue damage and parasite burden during chronic infection. Additionally, accumulation of reparative Treg cells in skeletal muscle was impaired in the absence of ST2. Together, these findings identify a tissue-adapted Treg cell population that integrates inflammatory and reparative programs to preserve skeletal muscle integrity during chronic parasitic infection.

**SUMMARY:** Chronic parasite infection drives the emergence of tissue-adapted regulatory T cells that integrate Th1-associated and tissue-repair programs. These cells accumulate in skeletal muscle, where they preserve tissue integrity while contributing to microbial control, highlighting the dynamic specialization of Treg cells across disease stages.

## INTRODUCTION

Foxp3⁺ CD4⁺ regulatory T (Treg) cells play essential roles in maintaining immune homeostasis and modulating immune responses across diverse settings. These roles rely on their capacity for functional specialization, whereby Treg cells adopt transcriptional programs guided by canonical Th cell–associated transcription factors (1). These programs confer migratory and functional properties tuned to local cues that enable context-dependent regulation of polarized effector T cell responses. Among the best characterized examples are Th1-like Treg cells, which acquire a transcriptional program that allows them to regulate type 1 inflammatory responses (2–4). This specialization is driven by the expression of the transcription factor T-bet, which coordinates the induction of genes associated with Th1 immunity while maintaining suppressive functions. A hallmark of Th1-like Treg cells is CXCR3, enabling colocalization within IFN-γ–rich sites and effective restraint of Th1/CD8⁺ responses during infection and autoimmunity (2, 4).

For many years, Treg cell research focused on lymphoid organs and circulation. More recent work has highlighted Treg cell populations in non-lymphoid tissues, revealing broader phenotypic and functional diversity. Tissue Treg cells exhibit substantial heterogeneity and have been proposed to play a primary role in tissue homeostasis together with immune suppression (5). Steady-state analyses indicate that tissue Treg cells form a dynamic pool shaped by migration and tissue residency programs (6). Within this category, “tissue-repair” Treg cells enriched in non-lymphoid sites promote tissue integrity and regeneration by engaging the IL-33/ST2 and amphiregulin (AREG) axis, a circuit that supports epithelial and stromal repair programs while remaining distinct from classical suppressive mechanisms (7–10). Local environmental cues further shape specialized Treg cell programs within individual tissues. For example, in lean visceral adipose tissue (VAT), ST2⁺ Treg cells support metabolic homeostasis by promoting insulin sensitivity and tissue remodeling, whereas CXCR3⁺ Treg cells restrain inflammatory responses and prevent glucose intolerance under high-fat diet conditions (11). In this setting, these populations have been described as mutually exclusive subsets that coexist within the same tissue and execute distinct, non-overlapping functions. Additional evidence for functional specialization comes from microbiota-dependent RORγ⁺ Treg cells, which differentiate in the gut and can migrate to sites of sterile tissue injury such as skeletal muscle (SM) and fatty liver, where they promote tissue repair and regulate local inflammation (12). These observations indicate that Treg cell specialization is associated with distinct transcriptional programs aligned with specific functional roles.

Infections represent highly complex scenarios where Treg cells must balance pathogen control with the prevention of excessive immune-mediated damage (13). Many pathogens, especially parasites, exploit Treg cell-mediated suppression to establish long-term persistence (14). Nevertheless, the role of Treg cells in chronic infections remains poorly understood, as these conditions are characterized by sustained localized inflammation despite reduced pathogen load, creating a setting in which Treg cells must adapt to persistent antigen exposure and ongoing tissue stress. Using *Trypanosoma cruzi* infection as a model of persistent parasitic infection, we have focused on defining the dynamics and functional specialization of Treg cells in this setting. Our previous studies demonstrated that, during the acute phase, Treg cells do not expand but become highly activated and acquire a Th1-like phenotype that suppresses CD8⁺ T cell immunity, at least in part through CD39-mediated mechanisms (15, 16).

Furthermore, we showed that the tissue-repair program is restrained during acute infection, correlating with extensive SM damage (17). While these findings highlighted a detrimental role for Treg cells in acute *T. cruzi* infection, clinical data from patients with chronic Chagas disease suggest an opposite trend: reduced frequency and/or impaired function of Treg cells are associated with more severe disease (18–20). This contrasting evidence raises the possibility that Treg cells undergo stage-specific reprogramming during infection, transitioning from predominantly Th1-like suppressive activity in acute disease to alternative functional states at chronicity. Whether chronic Treg cell responses reflect the sustained engagement of early effector-associated programs, the emergence of distinct tissue-adapted regulatory states, or a combination of both remains unclear.

To address this possibility, we characterized Treg cell responses during chronic *T. cruzi* infection and found that, while their numbers decline in the spleen, they accumulate in peripheral tissues, including SM, where they adopt a combined Th1 and tissue-repair program. Local depletion of these cells revealed their contribution to tissue homeostasis and parasite control. Moreover, perturbation of Treg cells during the acute phase of infection revealed that this stage contributes to shaping their long-term accumulation and functional properties, whereas IL-33/ST2 signaling emerged as a key pathway supporting their accumulation in SM. Together, our data indicate that chronic *T. cruzi* infection promotes the emergence of a specialized Treg cell subset with a key role in tissue homeostasis and parasite control, reflecting stage-specific adaptation of Treg cells to infection dynamics.

## RESULTS

### Chronic experimental *T. cruzi* infection recapitulates persistent parasitism, limited systemic inflammation, and skeletal muscle damage

To define the immunopathological landscape in which Treg cells operate during chronic *T. cruzi* infection, we first characterized tissue parasitism, immune infiltration, and organ damage. For that, Foxp3 reporter mice infected with 5,000 Tulahuen trypomastigotes were analyzed at ≥120 days post infection (dpi), a representative chronic time point (Fig. 1A).

**Figure 1.**
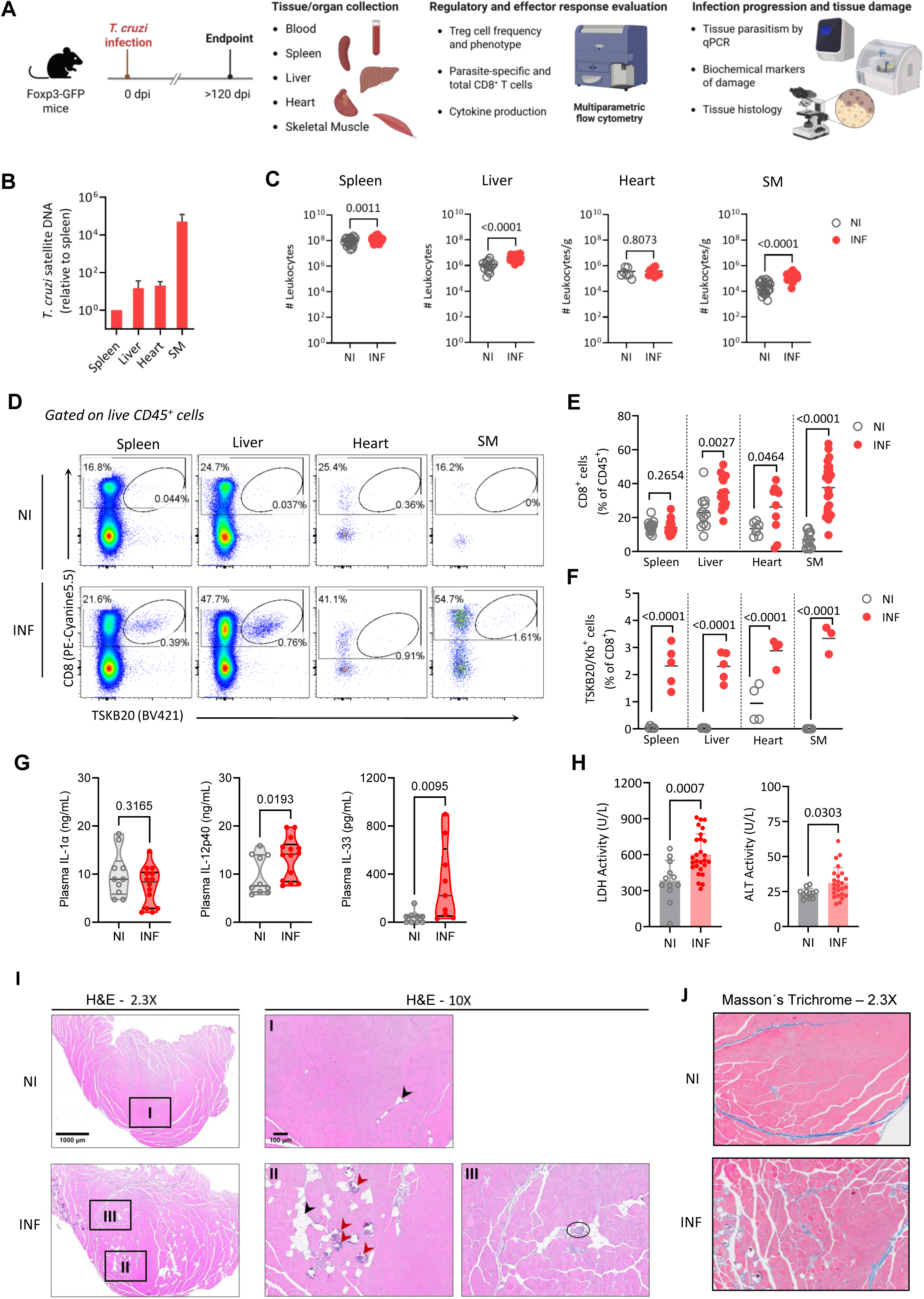
Chronic experimental *T. cruzi* infection is characterized by persistent parasitism, sustained immune infiltration, and skeletal muscle injury. (A) Experimental design. Foxp3-GFP mice were infected with 5,000 Tulahuen strain trypomastigotes (INF) or left non-infected (NI) and analyzed at ≥120 days post infection (dpi). Blood, spleen, liver, heart, and skeletal muscle (SM) were collected to assess immune response, parasite burden, and tissue damage. (B) Parasite burden in the indicated tissues from INF mice. (C) Total leukocyte counts. (D) Representative flow cytometry plots showing total and parasite-specific TSKB20/Kb^+^ CD8⁺ T cells within live CD45⁺ cells. (E) Frequency of total CD8⁺ T cells among CD45⁺ cells. (F) Frequency of parasite-specific TSKB20/Kb^+^ among CD8⁺ T cells. (G) Plasma levels of the indicated cytokines (N>9 mice/group). (H) Plasma biochemical markers of tissue damage. (I) Representative hematoxylin and eosin–stained sections of SM from NI and INF mice, showing necrotic fibers with dystrophic calcification (red arrow), adipose tissue accumulation (black arrow), and perivascular mononuclear infiltrates (black circle) in INF animals. Low- (2.3X) and high-magnification (10X) images are shown (N>4 mice/group). (J) Representative Masson’s trichrome–stained SM sections from NI and INF mice (magnification 10X), highlighting increased collagen deposition and interstitial fibrosis in INF animals (N>4 mice/group). Across panels, each symbol represents one mouse, bars indicate mean ± SD where applicable, and violin plots show the distribution of values within each group. Statistical analyses were performed as described in the Methods.

Quantitative PCR analysis detected *T. cruzi* satellite DNA in the spleen, liver, heart, and SM, with SM exhibiting the highest parasite burden (Fig. 1B). Concomitantly, chronically infected (INF) mice displayed increased splenocyte counts and enhanced leukocyte infiltration in the liver and SM, but not in the heart, compared to age-matched non-infected (NI) controls (Fig. 1C). These findings are consistent with sustained inflammation in the context of persistent parasitism.

We next evaluated the CD8⁺ T cell compartment (Fig. 1D-F) and found that INF mice exhibited similar frequencies of total CD8⁺ T cells in the spleen but significantly higher frequencies in the liver, heart, and SM (Fig. 1E). This increase translated into higher absolute numbers in the liver and SM, while numbers in the spleen and heart were comparable to NI mice (Suppl. Fig. S1A). Parasite-specific TSKB20/Kb⁺ CD8⁺ T cells were consistently detected in all tissues from INF animals at levels clearly above the background observed in NI mice (Fig. 1D and 1F; Suppl. Fig. S1B). Moreover, CD8⁺ T cells with an effector phenotype (CD44⁺CD62L⁻) were increased in both frequency and number across all target organs of INF mice compared to NI controls (Suppl. Fig. S1C-E). Together, these findings indicate that chronic infection is characterized by persistent parasite detection and sustained accumulation of effector CD8⁺ T cells into target tissues.

We next assessed systemic cytokine levels by quantifying more than 20 inflammatory mediators in plasma from NI and chronically INF mice. Only IL-1α, IL-12p40, and IL-33 were consistently detectable above the limit of detection across animals, and among them, only IL-12p40 and IL-33 were significantly increased in INF mice (Fig. 1G). Other cytokines were inconsistently detected in some NI or INF mice or remained below the limit of detection (Suppl. Fig. S1F), consistent with the absence of a broad systemic inflammatory cytokine signature at chronicity.

To assess tissue damage, we combined biochemical and histological analyses. INF mice displayed elevated lactate dehydrogenase (LDH) and alanine aminotransferase (ALT) activity (Fig. 1H), whereas aspartate aminotransferase (AST), creatine phosphokinase (CPK) and creatine phosphokinase of muscle and brain (CPK-MB) and glucose levels remained unchanged (Suppl. Fig. 1G). Given the muscular tropism of the Tulahuen strain, SM was selected for further analyses as a major site of parasite persistence, leukocyte infiltration, and tissue damage during chronic infection (17, 21). Histological analysis of SM confirmed the presence of necrosis and dystrophic calcification of muscle fibers, accompanied by perivascular mononuclear infiltrates, features absent in NI animals (Fig. 1I). In the same sections, an increased amount of adipose tissue was also evident in INF compared with NI mice. Masson’s trichrome staining further revealed enhanced collagen deposition and interstitial fibrosis in INF mice (Fig. 1J). Quantitative image analysis corroborated these findings, showing a higher proportion of SM area occupied by calcified, adipose, and fibrotic tissue in INF compared with NI mice, consistent with chronic tissue remodeling (Suppl. Fig. S1H).

Altogether, these data define the chronic phase of *T. cruzi* infection as a state of persistent parasitism associated with sustained presence of effector CD8⁺ T cells, limited systemic inflammatory cytokine responses, and moderate, organ-specific tissue damage, illustrating the immunopathological context of chronic infection.

### Treg cells undergo tissue-specific redistribution during chronic infection and accumulate in skeletal muscle

Having defined the chronic inflammatory environment, we next evaluated Treg cell distribution across tissues. We previously showed that Treg cell frequencies decline at the peak of acute *T. cruzi* infection in both lymphoid and non-lymphoid tissues (15, 17). To determine whether this pattern persists or changes at chronicity, we quantified Foxp3⁺ cells across tissues, including spleen, liver, heart, and SM, and show representative plots for spleen and SM (Fig. 2A). Consistent with acute infection, chronically INF mice exhibited reduced frequencies of Treg cells among splenic CD45⁺ cells compared with NI controls (Fig. 2B). the frequency of Treg cells among CD45⁺ cells was comparable in the liver and heart, whereas a marked increase was observed in SM3 (Fig. 2B). Analysis of absolute numbers revealed no significant changes in spleen and heart, while Treg cell counts were increased in liver and showed a pronounced expansion in SM (approximately threefold and tenfold, respectively) (Fig. 2C). These findings identify SM as a relevant site of Treg cell accumulation during chronic *T. cruzi* infection.

**Figure 2.**
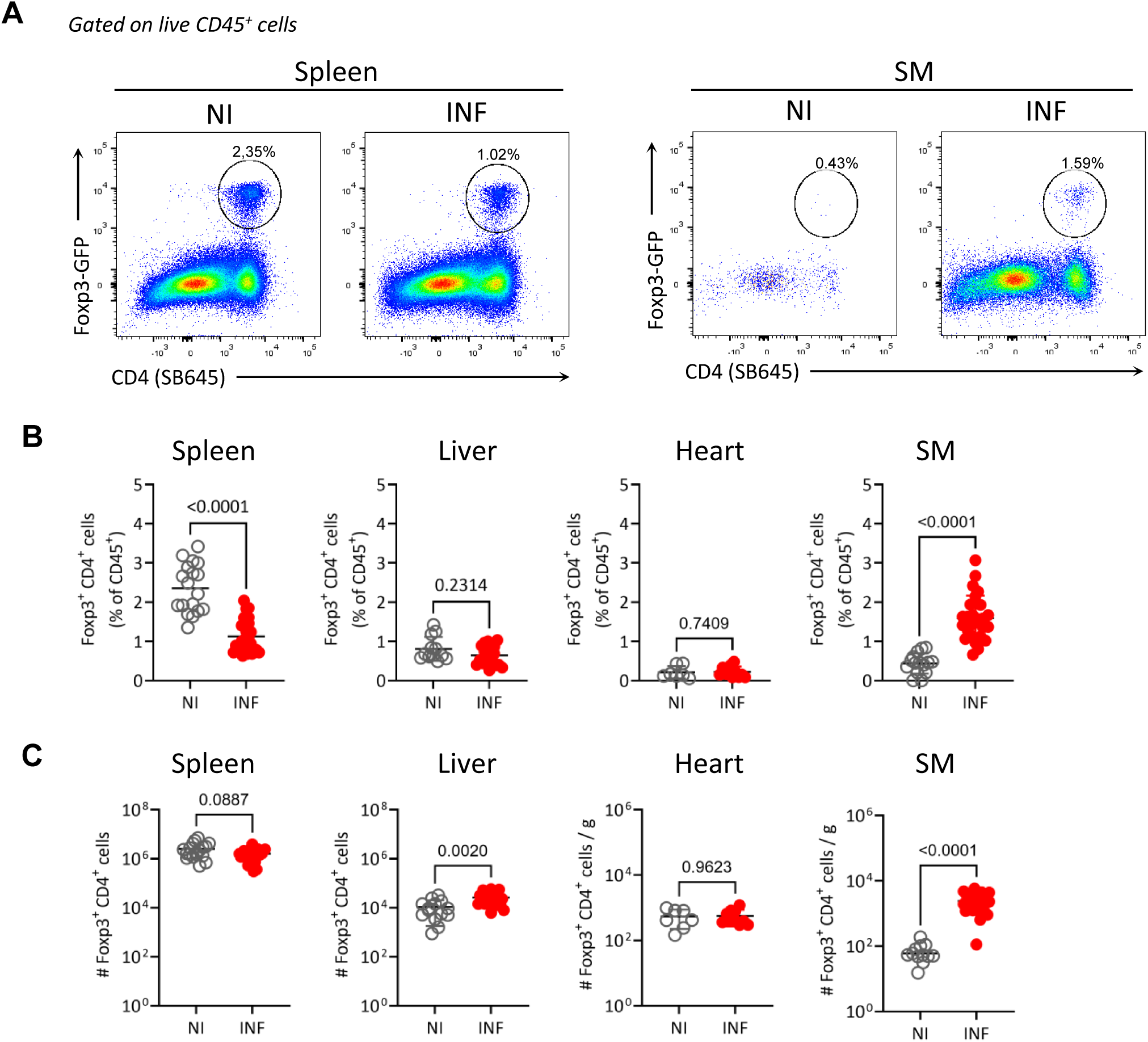
Regulatory T cells display tissue-specific dynamics and accumulate in skeletal muscle during chronic infection. (A) Representative flow cytometry plots showing Treg cells defined as Foxp3⁺ CD4^+^ cells within the CD45^+^ gate in spleen, and skeletal muscle (SM) from non-infected (NI) and chronically infected (INF) Foxp3-GFP mice at ≥120 days post infection. (B-C) Frequency among CD45⁺ cells (B) and absolute numbers (C) of Foxp3⁺ Treg cells in spleen, liver, heart, and SM. Across panels, each symbol represents one mouse. Statistical analyses were performed as described in the Methods.

### Muscle Treg cells are enriched for both Th1-like and tissue-repair signatures during chronic infection

Having identified that SM is a major site of parasite persistence, ongoing inflammation, and marked Treg cell accumulation during chronic *T. cruzi* infection, we next sought to investigate the features that distinguish these Tregs cells from those in lymphoid organs. To this end, we performed bulk RNA-seq of sorted Treg cells from SM and spleen of INF mice at ≥ 120 dpi (Suppl. Fig. S2A and S2B). Pearson correlation of normalized counts revealed a clear separation between SM and splenic Treg cells, with high similarity among samples within each group (Suppl. Fig. S2C). Using a threshold of shrunken log2FoldChange > 1 or < -1 and adjusted P < 0.05, gene expression analysis identified 398 differentially expressed genes between the two populations (Fig. 3A). SM Treg cells showed increased expression of genes associated with effector or tissue-adapted states, including *Id2* and *Prdm1*, whereas they expressed lower levels of transcripts linked to a lymphoid or central phenotype, such as *Ccr7, Sell, Tcf7, Lef1, Id3, Dtx1,* and *Bcl2* (10, 22, 23). Within the SM Treg cell population, we also observed upregulation of transcripts associated with tissue-repair functions, such as *Areg, Il1rl1, Klrg1, Gata3, Il10,* and *Cd274,* as well as increased expression of *Tbx21, Ifng, Tnf, Gzmb,* and *Cxcr3,* consistent with activated and Th1-like phenotype (12).

**Figure 3.**
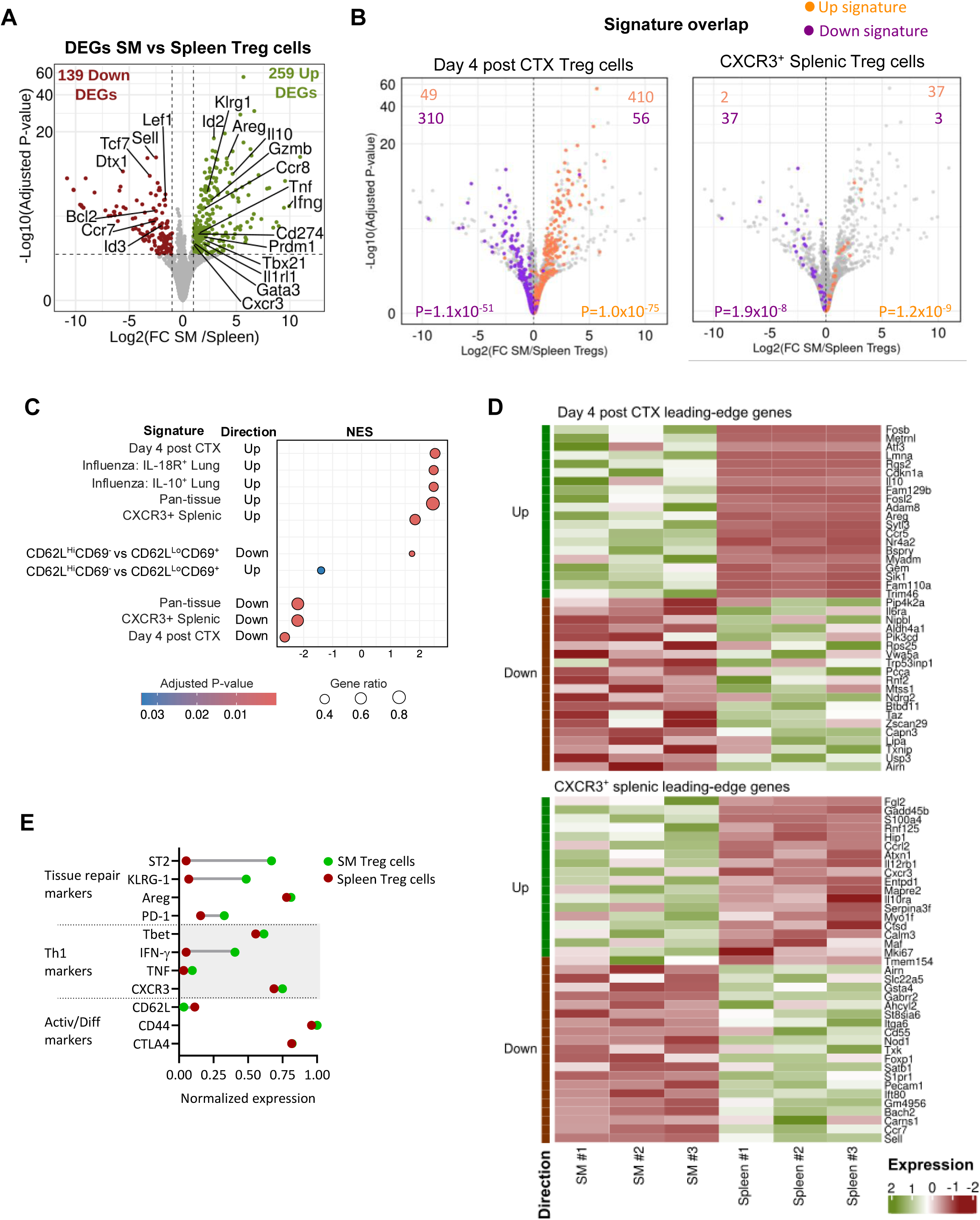
Skeletal muscle Treg cells are enriched for Th1-like and tissue-repair transcriptional signatures during chronic infection. (A) Volcano plot comparing gene expression between Treg cells isolated from skeletal muscle (SM) and spleen of chronically infected mice (≥120 dpi), highlighting genes significantly upregulated (Up DEGs, green) or downregulated (Down DEGs, red) in SM Treg cells. (B) Volcano plots showing differential gene expression between SM and splenic Treg cells. Genes belonging to published Treg cell signatures derived from day 4 post-CTX Treg cells (left) or CXCR3⁺ splenic Treg cells (right) are highlighted, with signature upregulated genes shown in orange and signature downregulated genes in purple. (C) Gene set enrichment analysis (GSEA) showing enrichment of selected transcriptional signatures in SM Treg cells. (D) Heat map showing scaled expression of leading-edge genes contributing to the enrichment of Day 4 post-CTX Treg cell and CXCR3⁺ splenic Treg cell signatures in SM versus spleen Treg cells. (E) Cleveland plot summarizing phenotypic differences between SM (green) and spleen Treg cells (brown), integrating markers associated with tissue repair, Th1 polarization, and activation and differentiation (N>4 mice/group).

To further interpret these transcriptional differences, we first compared SM Treg cells from chronically INF mice with published Treg cell signatures representative of distinct specialization states by testing the overlap of their predefined up- and downregulated gene sets. In this analysis, SM Treg cells showed a strong concordant overlap with the SM reparative Treg cell signature generated after cardiotoxin injury (7) (Fig. 3B), with a substantial proportion of genes upregulated in the reparative program also increased in SM Treg cells, and conversely for downregulated genes. In parallel, SM Treg cells also significantly overlapped with the signature of splenic CXCR3⁺ Th1-like Treg cells involved in pancreatic inflammation (24) (Fig. 3B). Additional signatures, including pan-tissue Treg cells (5), and IL-18R⁺ and IL-10⁺ Treg cell subsets arising during influenza infection (25), also showed significant overlap (Suppl. Fig. S2D-E).

We next performed gene set enrichment analysis (GSEA) using a genome-wide ranked list to evaluate enrichment against the same signatures and additional curated datasets related to Treg cell biology (Fig. 3C). Consistent with the overlap analysis, SM Treg cells were positively enriched for gene sets characteristic of Treg cells from injured or inflamed tissues, including pan-tissue, CXCR3-positive splenic, and IL-10- or IL-18R-positive lung populations, with reciprocal enrichment of some of the corresponding down signatures. Additionally, a GEO-derived dataset comparing naïve and effector Treg cells (GSE36527) supported an activated or effector-like state in SM Treg cells from chronically INF mice. To further illustrate the transcriptional programs underlying these enrichments, the top genes driving the enrichment from the injured tissue and CXCR3⁺ Th1-like Treg cell signatures are shown in Fig. 3D. These genes display higher expression in SM compared with splenic Treg cells, consistent with the enrichment of both tissue-repair and Th1-associated programs. Together, these analyses support the transcriptional enrichment of both Th1-like and tissue-repair signatures in SM Treg cells, consistent with an activated and tissue-adapted state.

To validate these transcriptional observations at the protein level, we performed phenotypic profiling of Treg cells from SM and spleen. The Cleveland plot in Fig. 3E summarizes the main differences observed between the two populations. SM Treg cells showed higher expression of molecules typically associated with tissue-repair responses, including ST2, KLRG1, AREG, and PD-1 (Suppl. Fig. S3A). In parallel, they displayed increased levels of Th1-associated markers, such as T-bet, IFN-γ, TNF, and CXCR3 (Suppl. Fig. S3B). Consistent with an effector phenotype, SM Treg cells exhibited reduced CD62L expression and increased CD44 expression compared with their splenic counterparts, whereas CTLA-4 levels were similar between the two populations (Suppl. Fig. S3C).

Together, these findings indicate that during chronic *T. cruzi* infection, Treg cells not only redistribute among tissues but also undergo tissue-specific specializations in SM, acquiring a transcriptional and phenotypic profile enriched for both Th1-like and tissue-repair features.

### Heterogeneity analysis reveals a dual-phenotype Treg cell subset enriched in skeletal muscle

Recent work has shown that ST2⁺ and CXCR3⁺ Treg cells can constitute distinct and reciprocally regulated populations in steady-state VAT (11). We therefore asked whether the dual transcriptional and phenotypic profile observed in SM Treg cells during chronic infection reflects discrete subsets or a single population displaying both features. To address this, we analyzed Treg cells from spleen, liver, and SM of chronically INF mice, with detailed phenotyping in SM. We first focused on Treg cells with a tissue-repair phenotype (trTreg cells), defined by coexpression of ST2 and KLRG1 (Fig. 4A). This subset showed increased frequency with comparable numbers in spleen, preserved frequency but increased absolute numbers in liver. In contrast, trTreg cells markedly accumulated in SM, both in frequency and absolute numbers, as this population was absent in NI mice (Fig. 4B, C).

**Figure 4.**
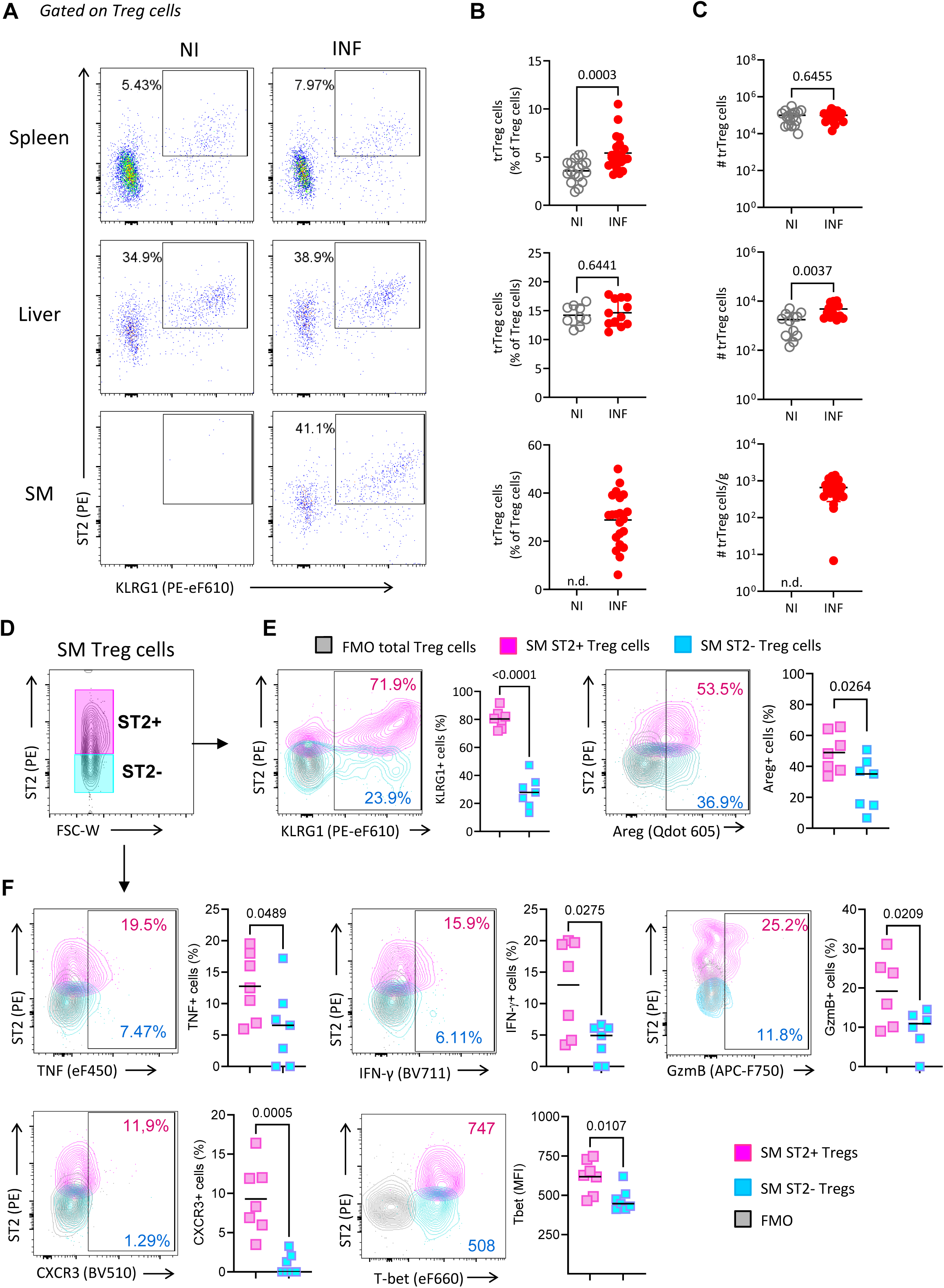
Heterogeneity analysis identifies an ST2⁺ Treg cell subset in skeletal muscle with combined tissue-repair and Th1-associated features during chronic infection. (A) Representative flow cytometry plots showing Treg cells with a tissue-repair phenotype, defined by coexpression of ST2 and KLRG1, in spleen, liver, and skeletal muscle (SM) from non-infected (NI) and chronically infected (INF) mice. (B) Frequency of ST2⁺ KLRG1⁺ Treg (trTreg) cells among total Treg cells according to A. (C) Absolute numbers of trTreg cells. (D) Gating strategy illustrating the classification of Treg cells into ST2⁺ and ST2⁻ populations in SM. (E) Representative flow cytometry plots and summary quantification showing the frequency of KLRG1⁺ and AREG⁺ cells within ST2⁺ and ST2⁻ Treg cell subsets in SM from INF mice. (F) Representative flow cytometry plots and summary quantification showing the expression of Th1-associated markers, including TNF, IFN-γ, granzyme B (GzmB), CXCR3, and T-bet, in ST2⁺ and ST2⁻ SM Treg cells from INF mice. Across panels, each symbol represents one mouse. Statistical analyses were performed as described in the Methods.

To further characterize the composition of the SM Treg cell compartment, we classified Treg cells according to ST2 expression into ST2⁺ and ST2⁻ populations (Fig. 4D) and compared additional markers linked to tissue repair or Th1 differentiation. As expected, ST2⁺ Treg cells displayed higher frequencies of KLRG1⁺ and AREG⁺ cells (Fig. 4E). Notably, ST2⁺ Treg cells from the SM of chronically INF mice exhibited higher frequencies of TNF⁺, IFN-γ⁺, and GzmB⁺ cells, along with increased CXCR3 and T-bet expression (MFI), compared with ST2⁻ Treg cells (Fig. 4F). Together, these data confirm the transcriptomic observations and demonstrate that during chronic *T. cruzi* infection, SM Treg cells represent a heterogeneous population enriched in ST2⁺ cells that coexpress tissue-repair and Th1-associated features at the cellular level.

To determine whether features associated with both Th1-like and tissue-repair programs are unique to chronic infection or also present at steady state, we analyzed the publicly available flow cytometry dataset FlowRepository FR-FCM-Z6ME from the Tissue Treg Project of the Liston–Dooley laboratory (6). Across eleven steady-state tissues (Fig. S4), all contained discernible ST2⁺ and ST2⁻ subsets, although their relative frequencies varied. ST2⁺ Treg cells were a minor population in blood, spleen, and inguinal lymph node (Suppl. Fig. S4A), more abundant in barrier tissues such as skin, female reproductive tract (FRT), and tongue (Suppl. Fig. S4B), and present at intermediate frequencies in metabolically active tissues including WAT, muscle, liver, heart, and kidney (Suppl. Fig. S4C). Of note, CXCR3⁺ cells were detectable within both ST2⁺ and ST2⁻ subsets across all tissues, with lower frequencies in lymphoid organs and variable representation across barrier and metabolically active tissues. When tissues were visualized according to these two parameters, circulatory/lymphoid, barrier, and metabolically active tissues occupied partially distinct regions of the ST2-CXCR3 space (Suppl. Fig. S4D). Overall, these datasets indicate that features associated with both tissue-repair and Th1 programs are present across most steady-state tissues, suggesting that they are not restricted to chronic infection.

### Transient systemic Treg cell depletion exerts limited effects during chronic infection

To assess the functional contribution of Treg cells during chronic infection, we used DEREG mice (Foxp3-DTR), which allow selective diphtheria toxin (DT)–mediated depletion of Foxp3⁺ Treg cells (26). Chronically INF DEREG mice received two consecutive intraperitoneal DT injections after 120 dpi, followed by blood sampling at 6 and 14 days after treatment (Tx+6d and Tx+14d, respectively) and tissue collection at Tx+14d (Suppl. Fig. S5A).

Treg cell depletion following systemic DT administration resulted in a marked reduction of circulating Foxp3⁺ CD4⁺ T cells at Tx+6d, which remained partially decreased at Tx+14d (Suppl. Fig. S5B). At Tx+14d, Treg cell frequencies were significantly reduced in the spleen, but not in SM (Suppl. Fig. S5C). No differences in the frequency of major immune cell populations, including B cells, CD4⁺ T cells, CD8⁺ T cells, monocytes, and neutrophils, were detected in spleen or SM (Suppl. Fig. S5D). Moreover, Treg cell depletion did not alter the frequency or absolute numbers of parasite-specific CD8⁺ T cells (Suppl. Fig. S5E). A deeper phenotypic analysis of activation markers and effector molecule expression on Foxp3^-^ CD4^+^ T (Tconv) and CD8⁺ T cells identified subtle changes, including increased frequencies of CD44⁺ CD8⁺ T cells and CD44⁺, CD25⁺, and CD107a⁺ Tconv cells in the spleen, as well as a reduction in the frequency of KLRG1⁺ CD8⁺ T cells in SM (Suppl. Fig. S5F). Evaluation of infection related parameters, including biochemical markers of tissue damage such as the activity of LDH, AST, ALT, CPK, and CPK-MB, as well as glycemia, revealed no significant impact of systemic Treg cell depletion (Suppl. Fig. S5G). Together, these results indicate that transient systemic DT-mediated Treg cell depletion after the establishment of chronic infection exerts only modest immunological effects.

### Local Treg cell depletion uncovers tissue-protective and antiparasitic functions in muscle

Because systemic DT administration did not efficiently eliminate Treg cells in SM and produced only limited and mostly systemic effects, we developed a protocol for local intramuscular Treg cell depletion. We performed intramuscular DT injections into three major posterior limb muscles (quadriceps, gastrocnemius and tibialis anterior) on both sides (Suppl. Fig. S6A). Control mice received intramuscular PBS injections into the same muscles, following the same schedule. Different DT doses were tested, and 1 ng/g was identified as the lowest dose that achieved effective Treg cell depletion in SM while inducing partial depletion of splenic Treg cells 48 h after treatment (Suppl. Fig. S6B).

Once an effective protocol for SM Treg cell depletion was established, we evaluated its impact on regulatory and effector responses, tissue damage, and parasite control at one and two weeks after DT injection (Fig. 5A). PBS-injected mice from both time points were pooled and analyzed as a single control group, as no major differences were observed between the day 7 and day 14 PBS groups. DT-treated mice were evaluated separately at Tx+7d and Tx+14d. We first evaluated the efficiency and persistence of Treg cell depletion. Treg cell frequencies in SM remained significantly reduced at both Tx+7d and Tx+14d, whereas splenic Treg cell frequencies showed only a trend toward reduction throughout the analysis period (Fig. 5B). Absolute cell counts confirmed a significant reduction in Treg cells in SM at both time points, while splenic Treg cell numbers were preserved (Fig. 5B). A similar pattern was observed for trTreg cells, which were decreased in SM but maintained in the spleen (Fig. 5C).

**Figure 5.**
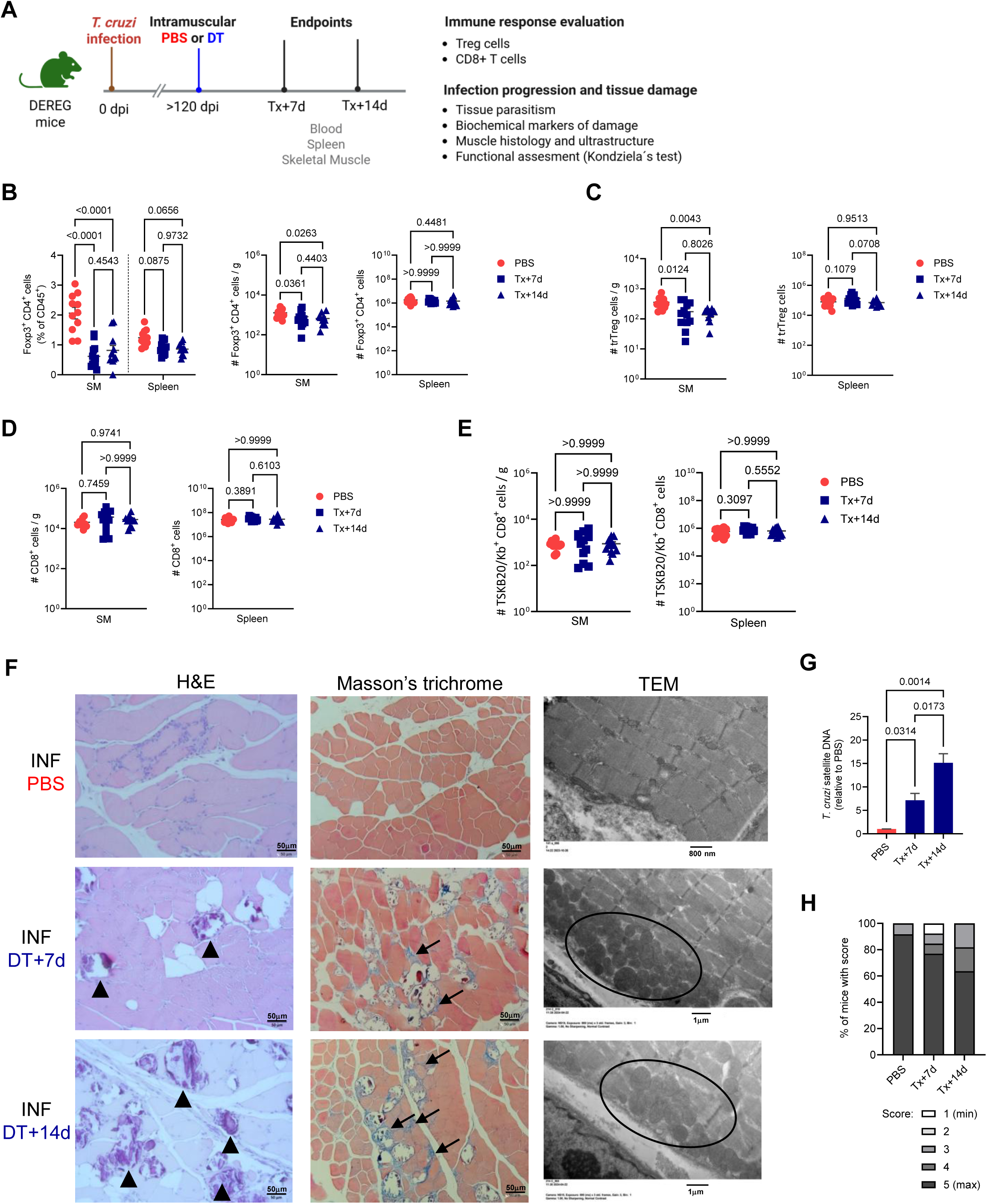
Local Treg cell depletion in skeletal muscle increases tissue damage and parasite burden during chronic infection. Chronically infected DEREG mice received intramuscular injections of PBS or diphtheria toxin (DT) into posterior limb muscles (quadriceps, gastrocnemius, and tibialis anterior) at ≥120 dpi. DT-treated mice were analyzed at Tx+7d and Tx+14d. PBS-treated mice from both time points were pooled and analyzed as a single control group. (A) Experimental scheme for local Treg cell depletion. (B) Frequency and absolute numbers of Foxp3⁺ Treg cells in skeletal muscle (SM) and spleen. (C) Absolute numbers of tissue-repair Treg (trTreg) cells in SM and spleen. (D-E) Absolute numbers of total (D) and parasite-specific TSKB20/Kb⁺ (E) CD8⁺ T cells in SM and spleen. (F) Representative SM sections stained with hematoxylin and eosin (H&E; left panels) showing areas of dystrophic calcification in DT-treated mice (arrowheads); Masson’s trichrome staining (middle panels) highlighting interstitial fibrosis, with delicate collagen tracts (blue stain) in DT-treated mice (arrows); and transmission electron microscopy images (TEM; right panels) showing subplasmalemmal mitochondrial accumulation in DT-treated animals (circles). Scale bars: H&E and Masson’s trichrome, 50 µm; TEM, PBS: 800 nm; DT: 1 µm. (G) Parasite burden in SM, determined by quantitative PCR for *T. cruzi* satellite DNA. (H) SM performance assessed by Kondziela’s inverted screen test. Across panels, each symbol represents one mouse. Bars indicate mean ± SD where applicable. Statistical analyses were performed as described in the Methods.

To determine whether local Treg depletion altered the inflammatory environment in SM, we evaluated effector CD8⁺ T cells, the composition of major immune populations, and local cytokine levels. No differences were detected in the number of total CD8⁺ T cells or parasite-specific CD8⁺ T cells in either SM or spleen at Tx+7d and Tx+14d (Fig. 5D and Fig. 5E). Likewise, the overall composition of major immune populations in SM was not significantly altered by local Treg depletion (Suppl. Fig. S7A). Only a subset of cytokines was consistently detected in SM lysates, including AREG, IFN-γ, IL-10, and IL-1β; however, no significant differences were observed between PBS- and DT-treated mice at either Tx+7d or Tx+14d (Suppl. Fig. S7B).

We next evaluated the impact of local Treg cell depletion on tissue damage and infection progression. No changes were observed in serum levels of AST, ALT, LDH, CPK, or glucose (Suppl. Fig. S7C). Histological examination of SM revealed substantially increased tissue damage in DT-treated chronically INF mice, characterized by extensive necrotic muscle fibers, prominent dystrophic calcification, and a multifocal mononuclear infiltrate involving interfascicular and perivascular regions (Fig. 5F, left panels). In contrast, PBS-treated mice displayed focal inflammatory infiltrates and limited areas of necrosis and calcification. Masson’s trichrome staining demonstrated mild interstitial fibrosis in both groups (Fig. 5F, middle panels), consistent with chronic remodeling. Necrotizing vasculitis was occasionally observed across samples. At the ultrastructural level, SM fibers from DT-treated mice exhibited marked myofibrillar disorganization, dilation of the smooth endoplasmic reticulum, and prominent subplasmalemmal mitochondrial accumulation (Fig. 5F, right panels), whereas PBS-treated mice preserved overall myofibrillar organization and lacked subplasmalemmal mitochondrial accumulation. These alterations were accompanied by increased tissue parasitism at both 7 and 14 days after treatment (Fig. 5G). Consistent with these findings, DT-treated mice showed a progressive decline in skeletal muscle performance, reflected by a reduced proportion of animals reaching the maximal score in Kondziela’s inverted screen test, a functional assay of muscle strength (Fig. 5H). To exclude non-specific effects of DT administration, we confirmed that DT injection in INF DEREG-negative littermates had no effect on SM Treg cell numbers (Suppl. Fig. S7D), other relevant immune populations such as CD8⁺ T cells (Suppl. Fig. S7E), muscle histology (Suppl. Fig. S7F), or performance in Kondziela’s test at 7 days post injection (Suppl. Fig. S7G).

Taken together, these findings identify a critical role for SM Treg cells in limiting tissue damage and controlling parasite burden during chronic infection.

### Early infection events and IL-33/ST2 signaling regulate long-term Treg cell accumulation in skeletal muscle

Given the relevance of SM Treg cells for tissue protection and parasite control, we next investigated how this population is established during infection. The acute phase is marked by strong Th1-like specialization of Treg cells, limited tissue-repair programming and effects on parasite control (15, 17), features that could influence their long-term accumulation. To test this possibility, we depleted Treg cells during the acute phase by administering DT at 5 and 6 dpi and analyzed mice at chronicity (≥120 dpi) (Fig. 6A).

**Figure 6.**
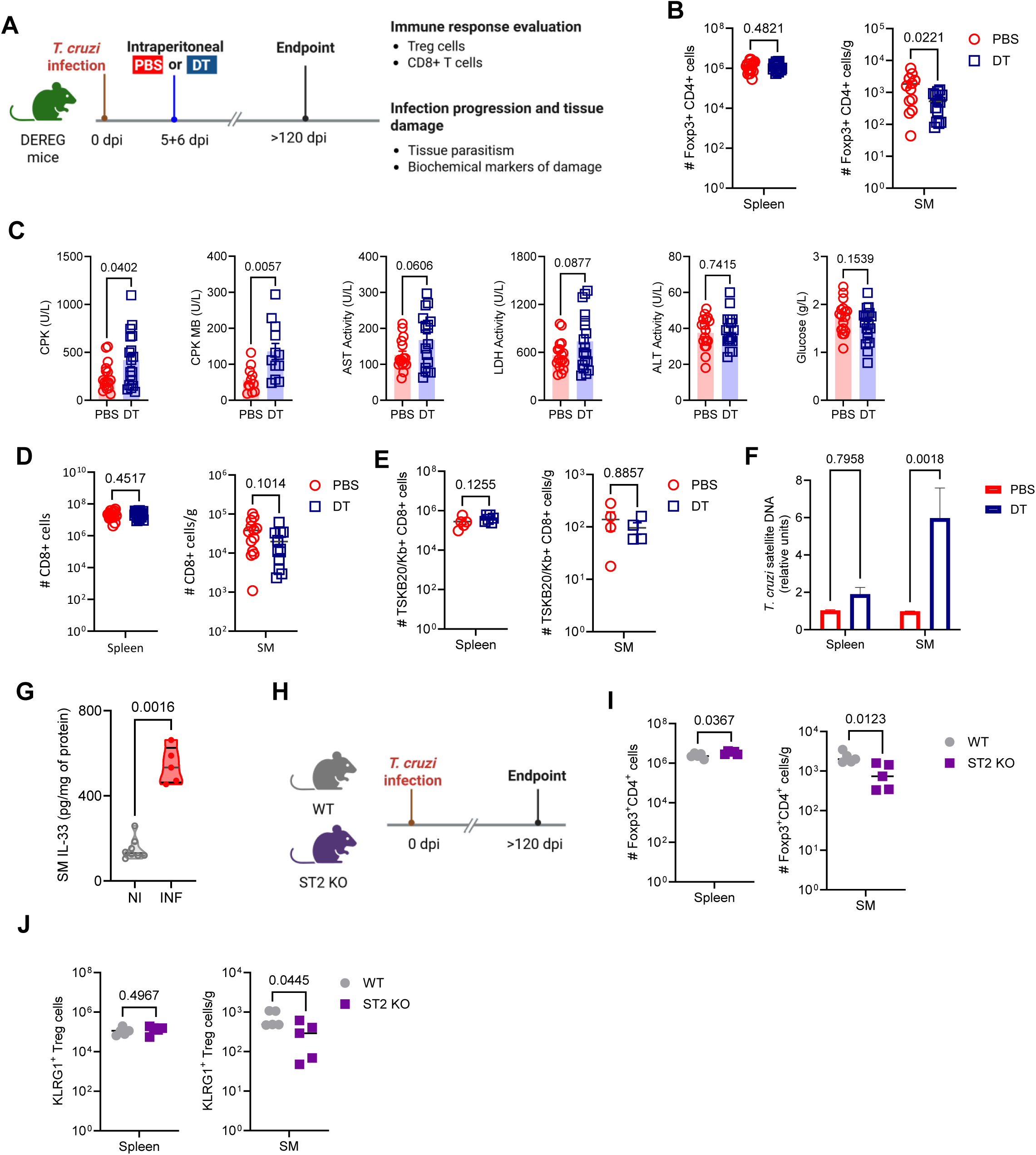
Signals acting during the acute phase shape long-term Treg cell accumulation in skeletal muscle. (A-F) DEREG mice were infected with *T. cruzi*, received diphtheria toxin (DT) at 5 and 6 days post infection (dpi), and were analyzed at the chronic stage (≥120 dpi). (A) Experimental scheme for acute-phase Treg cell depletion and chronic evaluation. (B) Absolute numbers of Foxp3⁺ Treg cells in spleen and skeletal muscle (SM). (C) Plasma biochemical markers of tissue damage. (D–E) Absolute numbers of total (D) and parasite-specific TSKB20/Kb⁺ (E) CD8⁺ T cells in spleen and SM. (F) Parasite burden determined by quantitative PCR for *T. cruzi* satellite DNA. (G) IL-33 concentration in SM lysates (N>5 mice/group). (H-J) Wild type (WT) and ST2-deficient mice (ST2 KO) were infected with *T. cruzi* and were evaluated at ≥120 dpi. (H) Experimental scheme. (I-J) Absolute numbers of total (I) and KLRG1⁺ (J) Foxp3^+^ Treg cells in spleen and SM. Across panels, each symbol represents one mouse. Bars indicate mean ± SD where applicable. Statistical analyses were performed as described in the Methods.

Quantification of Treg cells showed that acute DT administration had no long-term effect on splenic Treg cell numbers, yet it significantly reduced their accumulation in SM at chronicity (Fig. 6B). Acute-phase Treg cell depletion also resulted in increased biochemical markers of tissue damage at chronicity, including elevated CPK and CPK-MB activity and a trend toward higher AST and LDH activity, whereas ALT activity and glucose levels remained unchanged (Fig. 6C). Total and parasite-specific CD8⁺ T cell numbers were not affected in spleen or SM (Fig. 6D and Fig. 6E). Despite the lack of changes in CD8⁺ T cell immunity, the selective reduction in SM Treg cells was accompanied by increased parasite burden in SM, but not in spleen (Fig. 6F), consistent with the outcome of local Treg cell depletion during chronic infection.

These observations prompted us to test whether acute-phase signals shape the SM Treg cell compartment detectable at chronicity. To assess the contribution of acute IFN-γ signaling, we blocked IFN-γ during the acute phase and evaluated the long-term impact of this intervention on Treg cell populations (Fig. S8A). Acute IFN-γ blockade did not alter total Treg cell numbers in spleen or SM at chronicity (Fig. S8B). Consistent with our previous observations (17), IFN-γ blockade increased the size of the splenic trTreg cell population, whereas SM Treg cells remained unchanged (Fig. S8C). Likewise, biochemical markers of tissue damage and total or parasite-specific CD8⁺ T cell numbers were unaffected (Fig. S8D-F).

Because SM Treg cells display a prominent tissue-repair signature and express high levels of ST2, we investigated the potential contribution of IL-33/ST2 signaling. Analysis of cytokines in SM lysates revealed increased IL-33 levels in chronically infected mice compared with NI controls (Fig. 6G), whereas other cytokines were detected at lower levels and displayed greater inter-animal variability (Fig. S8G). We therefore studied the contribution of IL-33/ST2 signaling to SM Treg cell accumulation using chronically infected ST2-deficient mice (Fig. 6H). In these mice, Treg cell numbers were increased in spleen and reduced in SM (Fig. 6I). We next evaluated the KLRG1⁺ Treg cell subset, which was used as a proxy for reparative Treg cells because ST2 expression cannot be assessed in ST2-deficient hosts and KLRG1 and ST2 showed near-complete co-expression under WT conditions (Fig. 4A). Notably, KLRG1⁺ Treg cells were preserved in spleen but significantly reduced in SM (Fig. 6J). These findings indicate that IL-33/ST2 signaling is dispensable for maintaining splenic Treg cell populations but is required for the accumulation of muscle Treg cells, particularly a KLRG1⁺ subset associated with reparative functions.

Although our primary focus was on Treg cell accumulation, we also evaluated tissue damage and CD8⁺ T cell responses in ST2-deficient mice. ST2 deficiency did not induce major changes in biochemical markers of tissue damage, except for increased glycemia in ST2-deficient mice (Suppl. Fig. S8H). Total CD8⁺ T cell numbers were reduced in SM (Suppl. Fig. S8I), whereas no changes were detected in spleen or in parasite-specific CD8⁺ T cells in either tissue (Suppl. Fig. S8J).

Together, these results indicate that early infection events shape the long-term establishment of the SM Treg cell compartment and identify IL-33/ST2 signaling as a key pathway supporting its accumulation at chronicity.

### Cardiac tissue from chronic Chagas cardiomyopathy patients is enriched in Treg cell and Th1 specialized Treg cell transcriptional signatures

To explore whether features identified in murine SM Treg cells may relate to human disease, we turned to cardiac tissue from patients with chronic Chagas cardiomyopathy (CCC), the most clinically severe and extensively studied disease manifestation, and analyzed bulk RNA-seq data from left ventricular explants as reported in (27). In the original study, the authors described broad increases in immune-related signatures in CCC cardiac tissue compared with healthy donor controls and with patients with non-infectious dilated cardiomyopathy (DCM), consistent with substantial inflammatory infiltration dominated by Th1-type immune responses. Because bulk cardiac transcriptomes reflect a mixture of cardiomyocytes, stromal cells and infiltrating leukocytes, this approach does not resolve immune cell subsets. Nonetheless, coordinated variation in sets of Treg cell-related or specialization-associated transcripts may provide indirect insight into whether these programs are represented within the cardiac infiltrate. On this basis, we examined Treg cell core genes, as well as genes associated with Th1/Treg1 and reparative Treg cell programs, in CCC explants compared with healthy controls and DCM.

Plotting log₂ fold changes for CCC versus controls against CCC versus DCM suggested that many transcripts associated with Treg cells, as well as Th1/Treg1 and reparative programs, were higher in CCC cardiac tissue relative to both groups. These included canonical Treg-associated genes such as FOXP3, CTLA4, and IL10, as well as Th1/Treg1-associated transcripts such as *TBX21*, *IFNG*, and *CXCR3*. Transcripts associated with tissue-adapted Treg cell states, including *CCR8* and *BATF*, were also increased. In contrast, transcripts such as *IL2RA* and *IKZF2* (Helios) showed no clear change (Fig. 7A). Heat-map visualization was consistent with these trends, revealing coordinated expression of multiple Treg cell-associated transcripts in CCC cardiac tissue (Fig. 7B).

**Figure 7.**
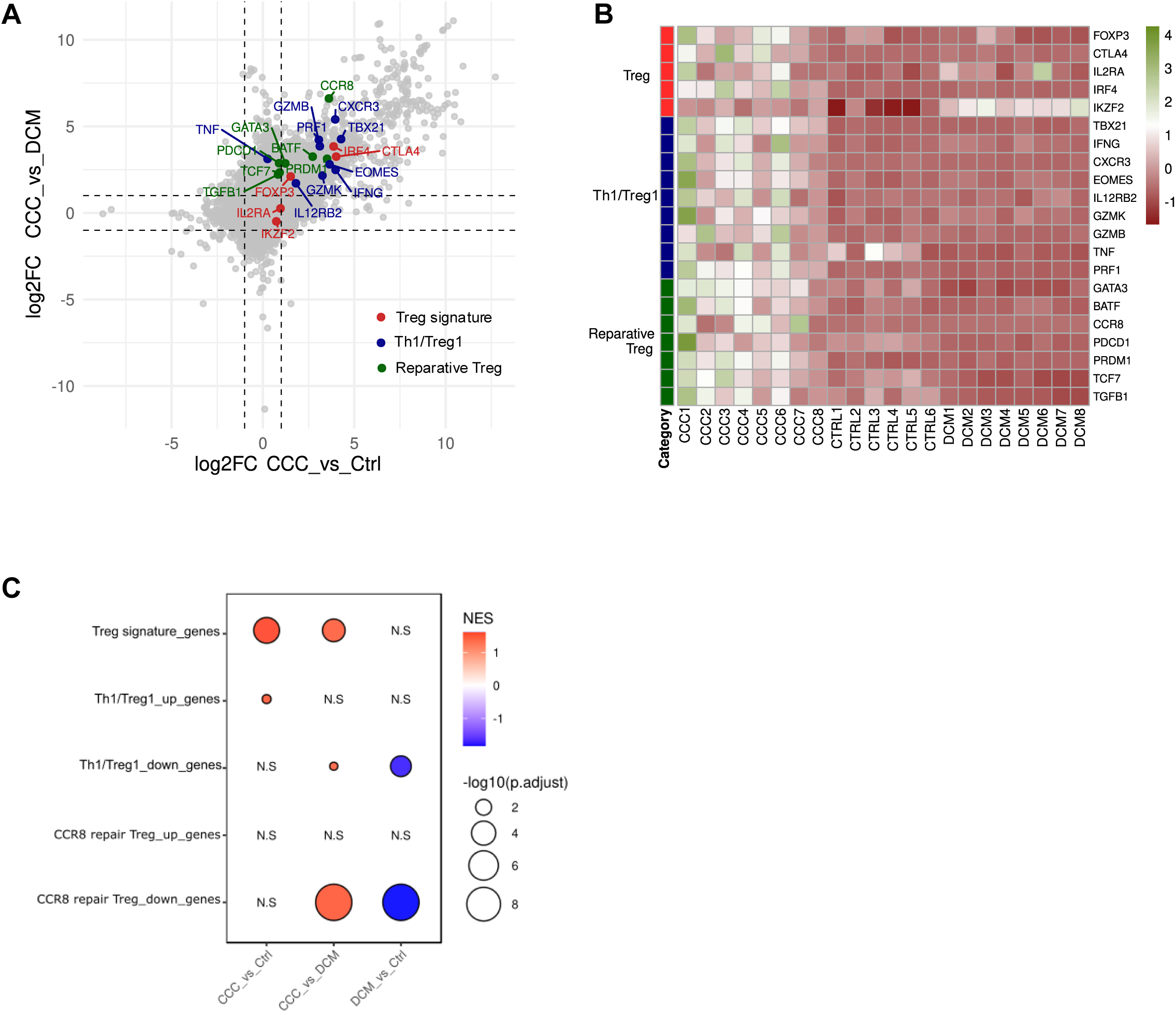
Cardiac tissue from chronic Chagas cardiomyopathy patients is enriched in Treg and Th1-specialized Treg cell transcriptional signatures. Publicly available bulk RNA-seq data from left ventricular explants of patients with chronic Chagas cardiomyopathy (CCC), non-infectious dilated cardiomyopathy (DCM), and healthy donor controls (Ctrl) (27) were reanalyzed in this study. (A) Scatter plot showing log₂ fold changes (FC) of all detected transcripts in cardiac tissue from patients with CCC compared with Ctrol (x-axis) and DCM (y-axis) patients. Genes belonging to Treg cell core, Th1-associated Treg (Treg1) cell, and reparative Treg cell signatures are highlighted and color-coded according to their assigned signature category. (B) Heat map showing relative expression of transcripts associated with Treg cell core, Treg1 cell, and reparative Treg cell signatures across cardiac samples from patients with CCC, DCM, and Ctrl. (C) Gene set enrichment analysis (GSEA) of CCC versus Ctrl, CCC versus DCM and DCM versus Ctrl comparisons using curated human Treg cell core, Treg1 cell, and reparative (CCR8⁺) Treg cell transcriptional signatures. Each column represents the indicated pairwise comparison. Dot color indicates the normalized enrichment score (NES), and dot size reflects –log₁₀ adjusted *P* value; non-significant comparisons are indicated as N.S.

We next performed GSEA using human Treg cell signatures and a Th1/Treg1-associated gene signature, originally defined in Treg1 cells by Hollbacher et al. and potentially reflecting broader Th1 responses in bulk tissue (28), as well as the CCR8⁺ reparative Treg cell signatures described by Delacher et al. (10) (Fig. 7C). The global Treg cell signature was positively enriched in CCC compared with both healthy donor controls and DCM, whereas no enrichment was detected in DCM relative to controls. CCC explants also showed positive enrichment of the Th1/Treg1-associated gene signature relative to controls and enrichment of the corresponding down signature relative to DCM. In contrast, reparative Treg cell programs did not show positive enrichment in any comparison; only the reparative down signature was enriched in CCC relative to DCM.

Despite the inherent limitations of bulk RNA-seq, these integrated signature analyses indicate that transcriptional programs associated with Treg cells and Th1-type immune responses are relatively enriched in CCC cardiac tissue, whereas reparative Treg cell programs are not prominently represented in this dataset.

## DISCUSSION

Infectious diseases impose evolving challenges on the immune system, requiring host responses to balance pathogen control with preservation of tissue integrity (29, 30). Treg cells are central to this balance and are increasingly recognized as a heterogeneous population shaped by tissue-derived cues and inflammatory context, rather than as a uniform suppressive lineage (5, 14). Here, we identify chronic *T. cruzi* infection as a model of tissue-specific regulatory reorganization in which Treg cells are preferentially enriched in SM, a site of parasite persistence (31–33), while remaining reduced in the spleen. These divergent compartmental dynamics indicate that regulatory programs are redefined according to tissue context and stage of infection.

In contrast to the predominantly Th1-aligned program described in splenic Treg cells during acute infection (15), SM Treg cells at chronicity display an integrated functional state combining type 1-associated and tissue-adaptive modules. Co-expression of CXCR3 and T-bet together with ST2 and AREG indicates that inflammatory and reparative programs can coexist within the same regulatory population under sustained inflammatory conditions. Although regulatory specialization at steady state can involve more segregated subsets (11), inflammatory environments appear to favor combinatorial transcriptional states. Similar concurrent expression of *Tbx21, Gzmb* and *Areg* transcripts has been described in Treg cells during acute mucosal viral infection (34), supporting the concept that inflammatory contexts can promote integrated regulatory programs rather than the expansion of a single dominant program. The functional relevance of this integrated state is underscored by the selective impact of local, but not systemic, Treg cell depletion during established chronic infection.

While transient systemic perturbation produced minimal changes, local removal in SM increased tissue damage and parasite burden, indicating that muscle Treg cells directly contribute to maintaining tissue integrity and restricting parasite expansion. In sterile injury models, Treg cells promote regeneration through AREG production and modulation of stromal compartments (7, 8). Our findings extend this framework to chronic infection, where Treg cells operate within a setting of persistent antigen exposure and ongoing low-grade inflammation.

Importantly, the increase in parasite load following local depletion occurred without detectable changes in inflammatory cytokine levels or in the number of parasite-specific CD8⁺ T cells or other major immune cell populations at chronic time points. These observations suggest that the relevant regulatory effects may not involve sustained alterations in bulk effector cell numbers. Instead, Treg cells may influence early activation dynamics, transient functional states, or cumulative tissue-protective processes that ultimately determine long-term parasite containment. In certain infection settings, Treg cells have been shown to promote the quality and durability of protective CD8⁺ T cell responses (35–37), supporting the possibility that temporally restricted regulatory effects contribute to pathogen control without necessarily altering steady-state cell numbers during established chronic infection. This contrasts with our previous observations during acute *T. cruzi* infection, where systemic Treg cell depletion enhanced parasite-specific CD8⁺ T cell responses by relieving CD39-dependent Treg suppression (16).

Treg cell depletion during the acute phase of infection further demonstrates that transient perturbation of regulatory dynamics during this stage has lasting consequences for both Treg cell accumulation and tissue integrity at chronicity, indicating that events occurring early after infection influence the establishment and long-term function of the chronic regulatory compartment. Although IFN-γ is a canonical inducer of T-bet expression in Treg cells (38, 39), IFN-γ blockade during the acute phase did not alter chronic Treg cell numbers in muscle, suggesting that IFN-γ alone may be insufficient to determine long-term Treg cell persistence in this tissue. In contrast, genetic disruption of ST2 revealed a non-redundant requirement for IL-33 signaling in maintaining the muscle Treg cell compartment, particularly the KLRG1⁺ subset associated with reparative features. Consistent with previous reports demonstrating a central role for IL-33 in stabilizing tissue-adapted Treg cell populations (5, 8, 40), these findings identify IL-33/ST2 as a key pathway supporting chronic Treg cell maintenance in SM during persistent infection.

Analysis of cardiac transcriptomic data from patients with chronic Chagas cardiomyopathy revealed enrichment of cell and Th1/Treg1-associated transcriptional signatures in diseased myocardium. Previous studies have reported differential regulatory profiles in peripheral blood of patients with indeterminate versus cardiac forms of Chagas disease (18, 19, 41, 42), although tissue-level data remain limited. Enrichment of a type 1-associated transcriptional signature in cardiac tissue, in the relative absence of a prominent reparative program, may reflect a regulatory state adapted to persistent inflammatory stress (43, 44), but with limited tissue-restorative capacity. While bulk RNA-seq does not resolve cellular heterogeneity, these findings are consistent with the hypothesis that imbalance between inflammatory and reparative regulatory modules may influence cardiac outcomes in chronic Chagas disease.

One limitation of this study is the absence of single-cell transcriptomic analysis of SM Treg cells, which would enable finer resolution of their heterogeneity and developmental trajectories. In addition, although we demonstrate that Treg cells restrain parasite burden in muscle, the precise mechanisms remain to be defined. Future studies should clarify whether regulatory cells act predominantly through modulation of tissue homeostasis, temporally restricted effects on effector responses, or other mechanisms that contribute to the balance between tissue integrity and parasite control during persistent infection.

Collectively, our data position chronic *T. cruzi* infection as a model of stage-dependent regulatory programming in which early infection events and IL-33/ST2 signaling contribute to the establishment of long-term tissue-adapted Treg cell states. The contrast between the Th1-skewed regulatory state observed during acute infection and the integrated inflammatory-reparative program that emerges at chronicity underscores the dynamic nature of Treg cell function across disease stages. Our data further demonstrate that early global interference with the regulatory compartment compromises the establishment of tissue-adapted Treg cell programs at later stages, with lasting consequences for tissue integrity and parasite control during chronic infection. Together, these findings indicate that therapeutic strategies aimed at modulating regulatory pathways during infection require temporal precision and careful consideration of stage- and tissue-specific Treg cell heterogeneity.

## MATERIALS AND METHODS

### Sex as a biological variable

Both male and female mice were included in all experiments. Animals were age matched (8 to 12 weeks old), randomly assigned to experimental groups, and data from both sexes were pooled for analysis because no sex-dependent differences were observed in the measured experimental outcomes. The findings are therefore expected to be applicable to both sexes.

### Mice

Wild-type C57BL/6 and BALB/c mice, Foxp3-GFP reporter mice (B6.Cg-Foxp3tm2Tch/J, RRID:IMSR_JAX:006772), DEREG mice (C57BL/6-Tg(Foxp3-DTR/EGFP)23.2Spar/Mmjax), and ST2-deficient mice on a C57BL/6 background were maintained as colonies at the Facultad de Ciencias Químicas, Universidad Nacional de Córdoba. Wild-type C57BL/6 and BALB/c founders were originally obtained from the Faculty of Veterinary Sciences, La Plata National University (La Plata, Argentina). Foxp3-GFP reporter and DEREG founders were originally purchased from The Jackson Laboratory (USA). ST2-deficient mice on a C57BL/6 background, generated by Dr Andrew McKenzie (45), were kindly provided by Dr Chad Steele (Tulane University). All animals were housed under a 12:12 h light-dark cycle with food and water *ad libitum*. Mice from different experimental groups were co-housed in the same cages. The institutional animal facility follows the recommendations of the Guide for the Care and Use of Experimental Animals published by the Canadian Council on Animal Care (CCAC). Histological and functional analyses were performed in a blinded manner when applicable.

### Study Approval

Mouse handling followed international ethical guidelines. All experimental procedures were conducted in compliance with the ethical standards set by the Institutional Animal Care and Use Committee of Facultad de Ciencias Químicas – Universidad Nacional de Córdoba, and were approved under protocol number RD-2022-2134-E-UNC-DEC#FCQ. Animal experiments are reported in accordance with the ARRIVE 2.0 guidelines.

### Key resources

Detailed information on antibodies, reagents, kits, tetramers, software, and other resources used in this study is provided in Supplemental Table 1.

### Parasites and experimental infection

For all experiments, the Tulahuen strain of *T. cruzi* was used. Bloodstream trypomastigotes were maintained in male BALB/c mice by serial passages every 10–11 days. For experimental infection, mice were intraperitoneally inoculated with 0.2 mL of PBS containing 5 × 10^3^ trypomastigotes obtained by dilution of blood from parasite passages. All infections were performed at similar times of the day. Unless otherwise indicated, mice were analyzed at ≥120 days post infection (dpi), a time point corresponding to a established chronic phase in this experimental model.

### Treg cell depletion

For systemic Treg cell depletion, DEREG mice were injected intraperitoneally with 25 ng of diphtheria toxin (DT) per gram of body weight (ng/g) diluted in PBS (16). DT was administered on two consecutive days at the indicated time points. PBS-injected DEREG mice were used as control. Treg cell ablation was confirmed in blood samples by flow cytometry two to five days after DT administration.

For local Treg cell depletion, unless otherwise stated, DEREG mice received 1 ng/g of DT by intramuscular injection in a total volume of 110 µL, distributed among both quadriceps, gastrocnemius, and tibialis anterior muscles (15–25 µL per muscle). This injection scheme was based on experimental frameworks originally established in cardiotoxin-induced SM injury models (7). DT was administered as a single dose at the indicated time point. To control for injection procedures and nonspecific effects of local DT administration, infected DEREG mice injected i.m. with PBS and infected DEREG-negative littermates receiving intramuscular DT injections were included. Treg cell depletion was evaluated in blood, spleen and SM samples by flow cytometry two days after DT administration. DT does not directly affect parasite viability, as previously reported (16).

### IFN-γ blockade

Mice received intraperitoneal injections with 200 µg of anti-IFN-γ (XMG1.2 clone) at the specified time points. Control mice received 200 µg of anti-horseradish peroxidase rat IgG1 (HRPN clone).

### Cell preparation

Blood was collected from the tail vein (20-50 μL) for Treg cell depletion control or for kinetic studies, while it was obtained via cardiac puncture (total blood) under anesthesia for endpoint experiments. In both cases, heparin was used as anticoagulant.

To obtain cell suspensions from solid tissues, euthanized mice were perfused with 10 mL cold Hanks’ Balanced Salt Solution. Spleens and livers were obtained and mashed through a tissue strainer. Liver infiltrating cells were obtained after 25 min centrifugation (600 g without brake) in a 35% and 67.5% bilayer Percoll gradient. The interphase containing leukocytes was recovered and washed. Erythrocytes in spleen and liver cell suspensions were lysed for 3 min using an ammonium chloride-potassium phosphate buffer (ACK Lysing Buffer). Heart and SM (quadriceps, gastrocnemius and tibialis anterior) were excised, minced and digested for 30 min in collagenase D (2 mg/mL, Roche) and DNase I (100 μg/mL, Roche). Digested tissues were filtered through a 70 μm filter and washed. Leukocytes were obtained after 25 min centrifugation (600 g without brake) in a 40% and 75% bilayer Percoll gradient. The interphase was recovered and washed. Cell numbers from all solid tissues were counted in Turk’s solution using a Neubauer chamber and used to calculate the number of specific subsets shown in several figures. For heart and SM samples, cell numbers were normalized to tissue weight and expressed as cells per gram of tissue.

### Parasite quantification

Parasitemia was assessed by counting the number of viable trypomastigotes in blood after lysis with a 0.87% ammonium chloride buffer. Abundance of *T. cruzi* satellite DNA in tissues was used to determine parasite burden. Genomic DNA was purified from 50 μg of tissue (SM, heart, liver and spleen) with TRIzol Reagent following manufacturer’s instructions. Satellite DNA from *T. cruzi* (GenBank AY520036) was quantified by real time PCR using a specific Custom Taqman Gene Expression Assay. Primers and probes sequences were previously described by Piron et al. (46). The samples were subjected to 45 PCR cycles in a thermocycler StepOnePlus Real-Time PCR System. Abundance of satellite DNA from *T. cruzi* was normalized to the abundance of GAPDH (Taqman Rodent GAPDH Control Reagent), quantified through the comparative ΔΔCT method and expressed as arbitrary units, as previously reported (15–17, 47, 48).

### Biochemical determinations

Plasma was collected after blood centrifugation for 8 min at 3000 rpm. Quantification of biochemical markers of tissue damage was performed at Laboratorio Biocon (Córdoba, Argentina) using a Dimension RXL Siemens analyzer. AST, ALT, LDH and CPK activity was determined by UV kinetic method, CPK-MB activity was evaluated by enzymatic method, while glucose concentration was assessed by kinetic/colorimetric method.

### Histological analysis

Perfused mouse quadriceps were fixed in 10% buffered formalin (pH 7), dehydrated, cleared, and embedded in paraffin. Sections (5 μm) were cut using a rotary microtome and mounted on glass slides. Hematoxylin and eosin (H&E) and Masson’s trichrome stainings were performed following standard procedures. Histopathological evaluation was performed by a pathologist under light microscopy. Images were acquired using a Phenoimager Fusion instrument (Akoya Biosciences) 2.3× and 10× magnification using a scientific CMOS camara and the Fusion software v1.0.8.

Quantitative image analysis was performed on whole-section images using ImageJ2 software. Adipose tissue, calcified areas, and fibrotic regions were identified based on morphological criteria in H&E- and Masson’s trichrome-stained sections and manually delineated. The percentage of tissue area occupied by each component was calculated relative to the total tissue area.

### Transmission electron microscopy

Perfused mouse quadriceps were fixed in Karnovsky’s fixative (4% formaldehyde and 2% glutaraldehyde in 0.1 M cacodylate buffer), post fixed with osmium tetroxide, dehydrated through graded acetones, and embedded in Spurr resin. Ultrathin sections were cut using an ultramicrotome and examined with a Hitachi HT7800 transmission electron microscope (Hitachi, Tokyo, Japan) operated at 100 kV, and images were acquired with an AMT NS15 digital camera (Advanced Microscopy Techniques Corp., Woburn, MA, USA) with system software v02.03. Ultrastructural analysis was performed by a trained pathologist.

### Flow cytometry

Single-cell suspensions obtained from blood and tissues were processed as described above. Antibodies used for flow cytometry are listed in Supplemental Table 1. For surface staining, cells were incubated with fluorochrome-conjugated antibodies in PBS containing 2% FBS for 20 min at 4 °C, including LIVE/DEAD Fixable Aqua Dead Cell Stain according to the manufacturer’s instructions. For the identification of *T. cruzi*–specific CD8⁺ T cells, cells were incubated with H-2Kᵇ TSKB20 (ANYKFTLV) tetramers for 20 min at 4 °C. For intracellular detection of transcription factors, cells were surface-stained, fixed, and permeabilized using the Foxp3/Transcription Factor Staining Buffer Set (Invitrogen) according to the manufacturer’s protocol. For intracellular cytokine detection, cells were stimulated for 2 h at 37 °C with PMA (50 ng/mL) and ionomycin (1 μg/mL) in the presence of brefeldin A and monensin, followed by surface staining, fixation, and permeabilization.

Samples were acquired on FACSCanto II or LSRFortessa flow cytometers. Data were analyzed using FlowJo software (version X.0.7). Doublets and dead cells were excluded based on forward scatter height versus area parameters and LIVE/DEAD staining, respectively. Regulatory T cells (Treg cells) were defined as CD45⁺ CD3⁺ CD4⁺ Foxp3/GFP⁺ cells. Fluorescence minus one (FMO) controls were used to define gating strategies for markers with continuous expression, and compensation was performed using single-stained compensation controls according to standard procedures.

### Analysis of public tissue Treg cell flow cytometry datasets

Publicly available flow cytometry datasets of tissue regulatory T cells were obtained from the Tissue Treg Project generated by the Liston–Dooley laboratory and deposited in FlowRepository (accession FR-FCM-Z6ME). These datasets are associated with the published study by Burton *et al.* describing the tissue-resident regulatory T cell pool across multiple non-lymphoid organs (6). Raw FCS files corresponding to Treg cells from multiple tissues were downloaded, including samples from both male and female mice. Files were concatenated by tissue of origin and subsequently reanalyzed using the same conventional gating strategy and analytical workflow applied to our own experimental samples. This approach ensured methodological consistency and enabled direct comparison of Treg cell phenotypic features across tissues under steady-state conditions.

### RNA-seq library preparation, sequencing and analysis

Treg cells were double-sorted using a BD FACSAria II Cell Sorter into TCL buffer containing 1% 2-mercaptoethanol (2-ME), using the strategy depicted in Supplemental Figure S2B. For transcriptomic analyses, three independent pooled samples per tissue (spleen and SM) were generated from 4–6 infected mice. Within each biological replicate, spleen and SM Treg cells were obtained from the same animals and sorted independently. Libraries were constructed, sequenced, and the data were processed according to the ImmGen protocol (https://www.immgen.org/img/Protocols/ImmGenULI_RNAseq_methods.pdf), and as previously described (49). The Smartseq2 protocol was used for library construction (50, 51), and RNAClean XP beads were used to capture and purify RNA. An anchored oligo(dT) primer (59–AAGCAGTGGTATCAACGCAGAGTACT30VN-39) was used to select poly-A mRNA, which was subsequently reverse-transcribed to cDNA and PCR amplified. The Nextera XT DNA Library Preparation Kit was used to perform Tn5 transposon-based fragmentation. Barcoded primers were used in another round of PCR amplification (12 cycles) to introduce unique combinations of Illumina P5 and P7 barcodes to different samples, enabling pooling of samples before sequencing. Sequencing was performed on an Illumina NextSeq500 (two full NextSeq runs per batch of 96 samples, for approximately 10 million raw reads per sample on average) using 2x 38-bp reads with no further trimming. STAR 2.7.3a was used to align the sequencing reads to the mouse genome (GENCODE GRCm38/mm10 primary assembly and gene annotations vM25). Transcripts annotated as ribosomal RNA were removed. Quantification at the gene level was calculated using the Subread 2.0 command featureCounts.

For the analysis of the transcriptome of CCC heart explants, the raw counts matrix of the human heart-explant RNAseq dataset from (27) was downloaded from Gene Expression Omnibus (Accession number GSE191081).

The number of reads per sample was checked, and in both cases all samples contained an appropriate number of reads and were included in the analysis. Genes which did not have more than 10 reads in at least two samples were filtered out. Normalization and expression comparison were performed in the Bioconductor package DESeq2 (version 1.48.2, (52). Fold changes were shrunk using the apeglm algorithm implemented in DESeq2 (53), and the resulting shrunken fold changes were used for all subsequent analyses. Genes were defined as differentially expressed if their shrunken log2(FoldChange) was above 1 or below -1, and adjusted P-value was below 0.05. For the human dataset, ensembl gene identifiers were converted to gene symbols using the biomaRt package in R (54). The normalized counts, raw and shrunken log2(FoldChange), p-values and adjusted p-values are included in Supplemental Table 2 (mouse spleen vs SM Treg cells dataset) and Supplemental Table 3 (Heart explant dataset).

To analyze signatures by fisher’s exact test, the number of genes with positive and negative Log2(FoldChange) found and not found in the signature were used to build a contingency table. Fisher’s exact test was used on these tables to calculate P-values, which were adjusted for multiple comparisons using the Benjamini-Hochberg method.

Gene set enrichment analysis (GSEA) was performed using package clusterProfiler (version 4.16.0, (55) and some cell signatures were accessed from the Molecular Signature Database (MSigDB) using msigdbr (version 25.1.1), or derived from publicly available GEO datasets.

Expression heatmaps show z-scores based on the normalized counts. Plots were generated in R using packages clusterProfiler, ggplot2, pheatmap and complexheatmap.

The RNA-seq data generated in this study have been deposited in the NCBÍs Gene Expression Omnibus (GEO) (56) and will be released upon publication.

### Kondziela’s inverted screen test

Strength test was performed according to the protocol reported by Robert M. J. Deacon (57). Briefly, mice were kept in the experimental room for 20 minutes before the test to ensure proper adaptation to the environment. Each mouse was placed in the center of a 43 cm^2^ wire mesh consisting of 12 mm squares of 1 mm diameter wire and surrounded by a 4 cm deep wooden frame. Screen was inverted and held steadily 40-50 cm above the bench and time was measured until the mouse fell off. A score was assigned to each animal according to the following criteria: falling between 1-10 sec = 1; falling between 11-25 sec = 2; falling between 26-60 sec = 3; falling between 61-90 sec = 4; falling after 90 sec = 5. Maximum test duration was 2 minutes.

### Cytokine quantification

Plasma and SM samples were processed as described above. Plasma samples were analyzed without further dilution unless otherwise indicated. SM lysates were prepared by homogenization in PBS containing 0.5% BSA, 0.4 M NaCl, 1 mM EDTA, 0.05% Tween 20, and a protease inhibitor cocktail (Roche), followed by centrifugation at 10,000 g for 10 min, adapted from (58). For all cytokine determinations performed in SM, values were normalized to total protein content measured by the Bradford assay (Bio-Rad).

IL-33 concentrations were determined using a Mouse IL-33 ELISA kit. Calibration curves were generated using GraphPad Prism software (version 9.0). Absorbance was measured at 450 nm (ELISA) and 595 nm (protein quantification) using a Synergy HT multi-mode microplate reader (BioTek). Other cytokine concentrations in plasma and SM samples were quantified using the LEGENDplex Th Cytokine Panel (IFN-γ, IL-5, TNF, IL-2, IL-6, IL-4, IL-10, IL-9, IL-17A, IL17F, IL-22, IL-13), Cytokine Panel 2 (IL-12p70, IL-1α, IL-1β, IL-3, IL-12p40, IL-23, IL-7, IL-11, IL-27, IFN-β, GM-CSF, TSLP) or Custom bead-based multiplex assays according to the manufacturer’s instructions. Data were acquired by flow cytometry, and cytokine concentrations were calculated using the LEGENDplex Data Analysis Software based on standard curves generated for each analyte.

### Statistics

Descriptive statistics were calculated for each experimental group. Normality of data distribution was assessed using the Shapiro–Wilk test. Statistical significance was determined using two-tailed *t* tests or one-way ANOVA for normally distributed data, and two-tailed Mann–Whitney or Kruskal–Wallis tests for non-normally distributed data, as appropriate. *P* values < 0.05 were considered statistically significant, whereas *P* values < 0.1 were considered a trend. Outliers were identified using the ROUT method. All statistical analyses and graphical representations were performed using GraphPad Prism version 10.0. Data are presented as mean ± SD. The experimental unit was an individual mouse unless otherwise indicated.

Animals or samples were excluded only due to technical issues affecting sample quality or analysis (e.g., tissue contamination during dissection or sample loss during processing). Sample sizes were selected based on previous studies using the same experimental model and on our prior experience detecting biologically relevant differences in immune and pathological parameters. Most datasets were obtained from multiple independent experiments, as the calculated sample size required to achieve statistical significance was distributed across one to three independent assays. In cases where data are shown from a representative experiment, comparable statistically significant differences were consistently reproduced across independent experiments. The number of animals included in each experimental group is indicated in the dot plots, unless otherwise specified.

### AI Language Model Assistance

ChatGPT (OpenAI; GPT-4 and GPT-5 series models available through the ChatGPT platform) was used to assist in editing portions of the manuscript text. All generated content was reviewed and revised by the authors, who take full responsibility for the final manuscript.

## AUTHOR CONTRIBUTIONS

CLAF, SB, CMSG, and EVAR designed the study. CLAF, SB, CMSG, YG, JG, and CR performed experiments and acquired data. CLAF, SB, CMSG, and EVAR analyzed and interpreted the data. BSH contributed to RNA-seq data analysis and interpretation. JHM performed histopathological analyses. MCAV, AG and CLM contributed to data interpretation and critical discussion. EVAR conceived and supervised the study and wrote the manuscript. All authors reviewed, edited, and approved the final manuscript.

CLAF, SB, and CMSG contributed equally as first authors. Authorship order among the co-first authors was determined alphabetically.

## FUNDING SUPPORT

This work was supported by the Secretaría de Ciencia y Técnica–Universidad Nacional de Córdoba under Grant 33620230100016CB, Agencia Nacional de Promoción de la Investigación, el Desarrollo Tecnológico y la Innovación under Award Prestamo BID PICT 2020 SERIE A – 0487 and the National Institute of Allergy and Infectious Diseases of the National Institutes of Health under Award Number R01AI169482. The content is solely the responsibility of the authors and does not necessarily represent the official views of the funding agencies.

## AKNOWLEDGEMENTS

We thank P. A. Abadie, P. M. Crespo, V. Blanco, D. Lutti, C. Noriega, F. A. Frontera, S. R. Oms, R. E. Villarreal, G. Furlán, N. M. Maldonado, A. Romero, L. V. Gatica, M. S. Miró, D. A. Paira and L. Reyna (Centro de Investigaciones en Bioquímica Clínica e Inmunología), as well as A. V. Juárez and C. Leimgruber (Instituto de Investigaciones en Ciencias de la Salud) for their excellent technical assistance. We acknowledge the NIH Tetramer Core Facility for provision of TSKB20 tetramers. We thank Ayelén González Montoro (ORCID 0000-0002-6978-8284, HighPeak Bioinformatics) for her expertise and support in bioinformatic analyses.

## Supplemental Figure Legends

**Supplemental Figure S1.**
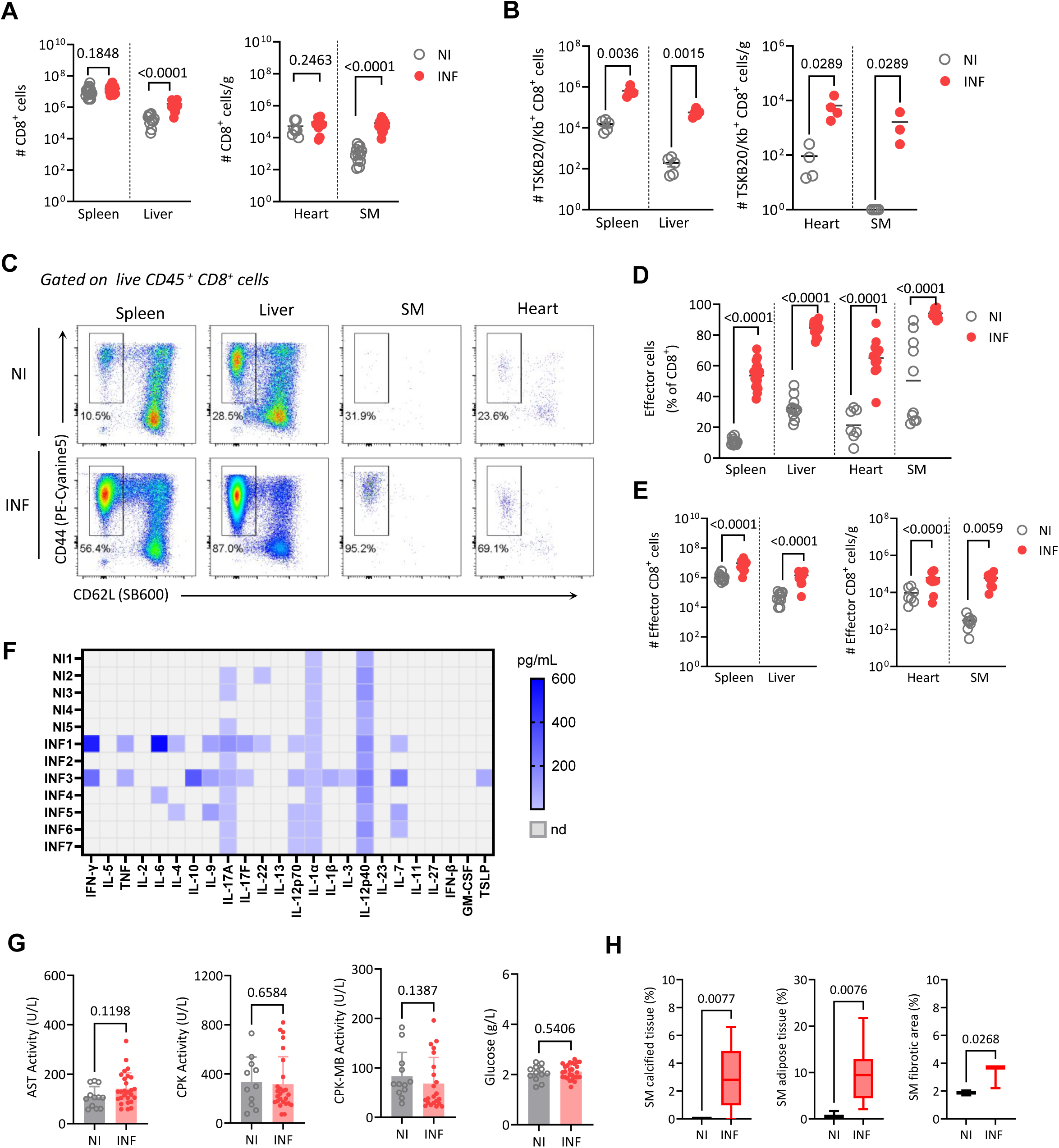
Chronic *T. cruzi* infection induces sustained immune activation and tissue alterations across target organs. Foxp3-GFP mice were infected with 5,000 Tulahuen strain trypomastigotes (INF) or left non-infected (NI) and analyzed at ≥120 days post infection (dpi). Blood, spleen, liver, heart, and skeletal muscle (SM) were collected to assess immune response, parasite burden, and tissue damage. (A-B) Absolute numbers of total (A) and parasite-specific TSKB20/Kb^+^ (B) CD8⁺ T cells. (C) Representative flow cytometry showing effector CD8⁺ T cells (CD44⁺CD62L⁻) within CD45⁺ CD8⁺ cells. (D) Frequency of effector CD8⁺ T cells among total CD8⁺ T cells. (E) Absolute numbers of effector CD8⁺ T cells. (F) Heat map showing relative levels of cytokines detected in plasma from NI and INF mice. Each row represents one mouse. (G) Plasma biochemical markers of tissue damage and glycemic status. (H) Quantification of tissue areas corresponding to calcification, adipose tissue, and fibrosis in SM sections. Across panels, each symbol represents one mouse, bars indicate mean ± SD where applicable. Boxes indicate the median with interquartile range, and whiskers denote minimum to maximum values. Statistical analyses were performed as described in the Methods.

**Supplemental Figure S2.**
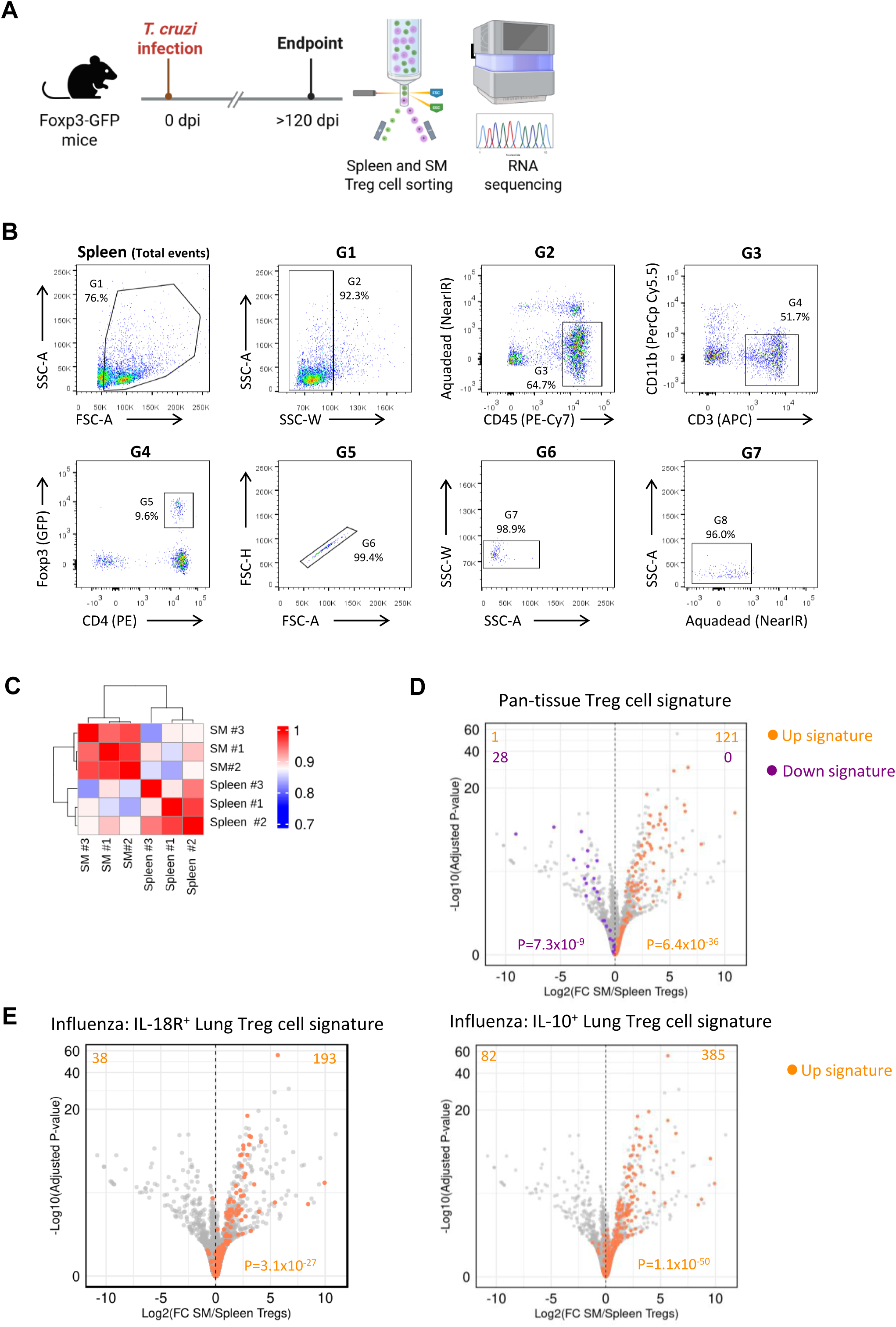
Experimental workflow and transcriptional analysis of skeletal muscle and splenic Treg cells. (A) Experimental scheme for transcriptional profiling of splenic and skeletal muscle (SM) Treg cells from chronically infected mice. (B) Sorting strategy used to purify Foxp3⁺ Treg cells, exemplified in the spleen. (C) Pearson correlation matrix of normalized RNA-seq counts showing clustering of samples according to tissue of origin. Color scale indicates Pearson correlation coefficients (r). (D and E) Volcano plots showing differential gene expression between SM and splenic Treg cells. Genes belonging to selected published Treg cell transcriptional signatures are highlighted, including pan-tissue, and influenza-associated IL-18R⁺ and IL-10⁺ lung Treg cell signatures, with signature upregulated genes shown in orange and signature downregulated genes in purple.

**Supplemental Figure S3.**
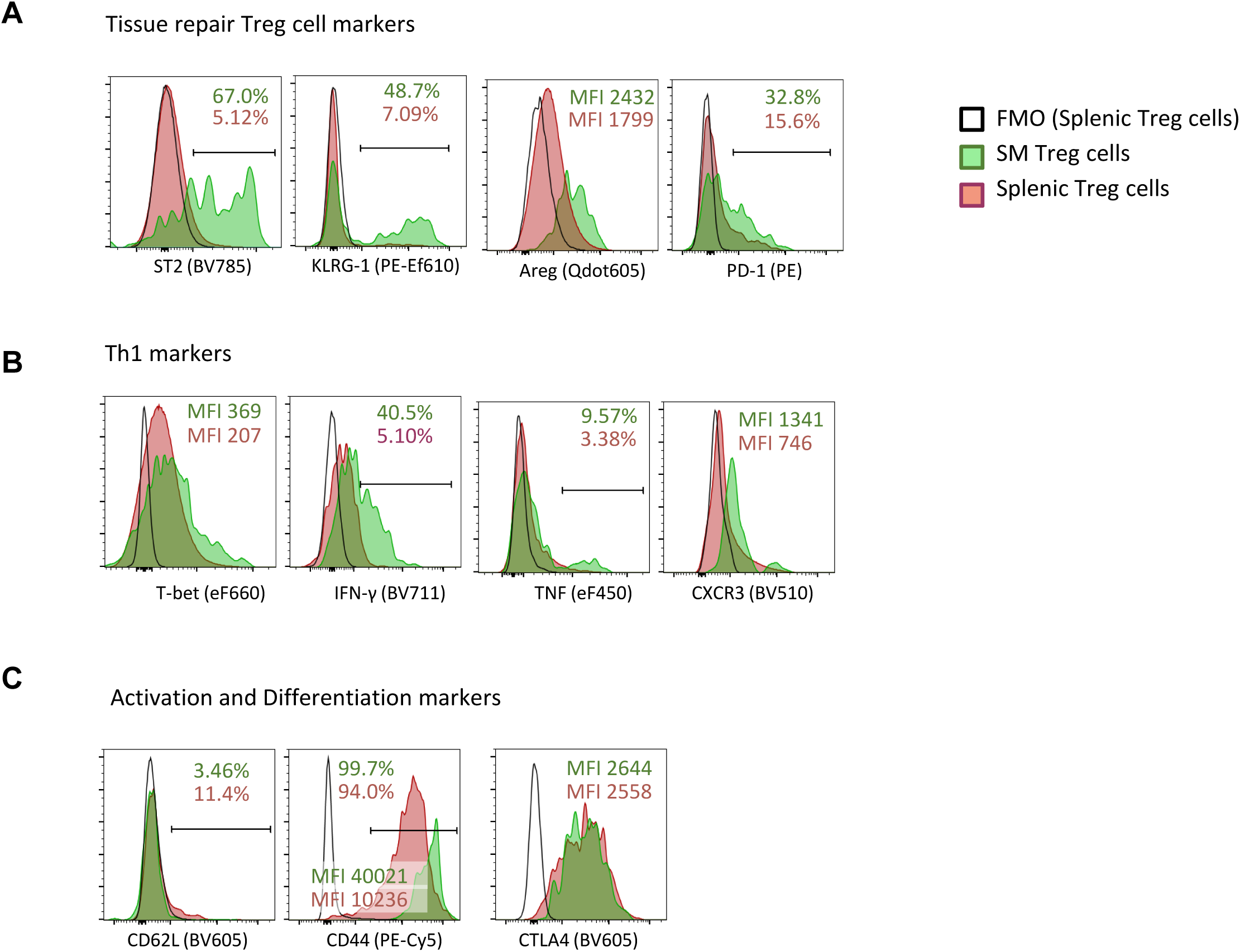
Phenotypic validation of Th1-like and tissue-repair features in skeletal muscle Treg cells. (A-C) Representative histograms showing expression of tissue-repair–associated markers (A), Th1-associated transcription factors and cytokines (B), and activation and differentiation markers (C) in skeletal muscle (SM) and splenic Treg cells from chronically infected mice. Percentages or mean fluorescence intensity (MFI) values are indicated within each panel. Data are representative of chronically INF mice (N≥4 per group) analyzed at ≥ 120 days post infection. Fluorescence minus one (FMO) controls were performed using splenic Treg cells.

**Supplemental Figure S4.**
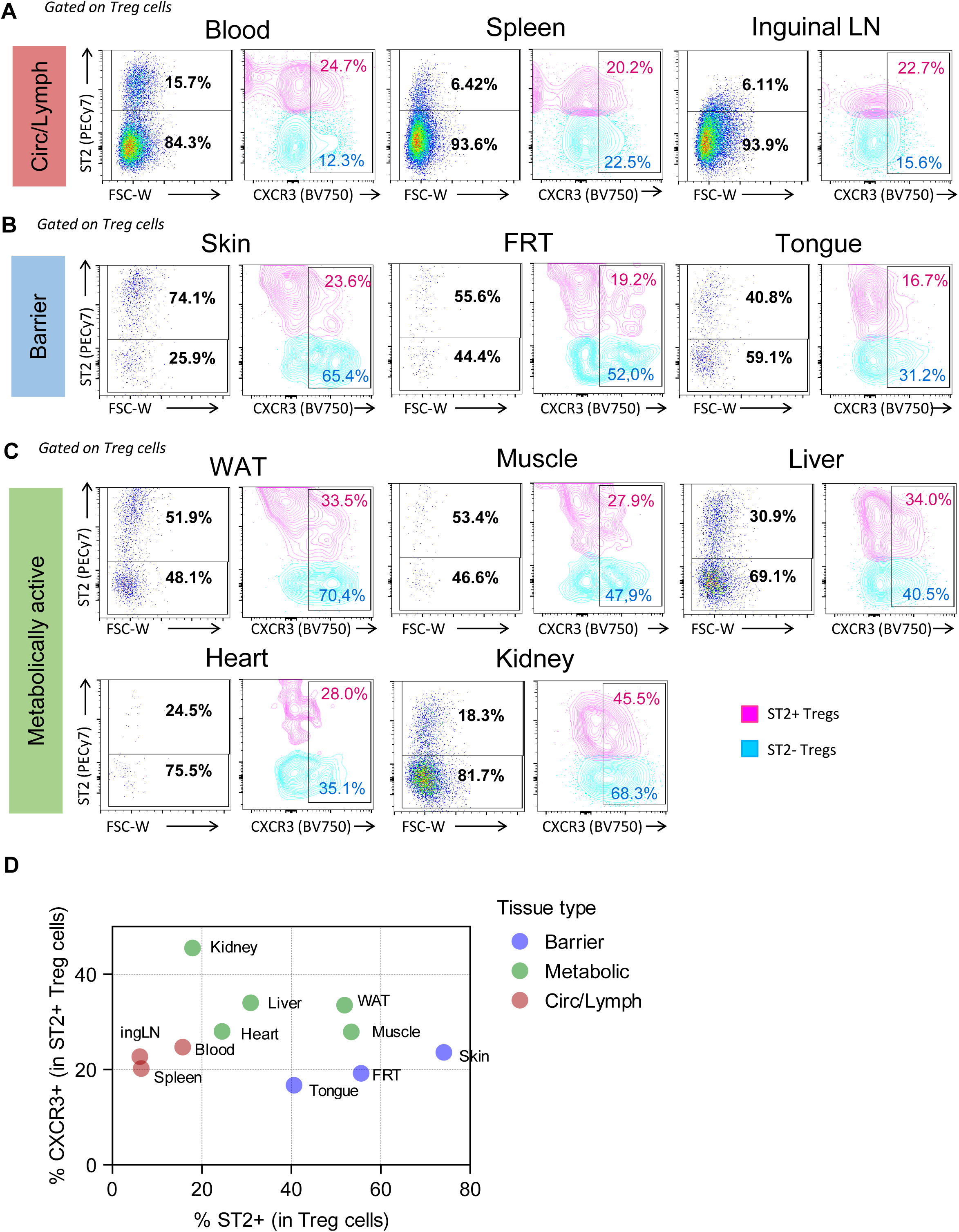
ST2⁺ and CXCR3⁺ Treg cell populations coexist across tissues in mice under steady-state conditions. Flow cytometry data were obtained from the Tissue Treg Project (FlowRepository FR-FCM-Z6ME) and analyzed in this study. (A–C) Representative flow cytometry plots showing ST2 and CXCR3 expression among Treg cells across tissue categories: circulating or lymphoid tissues (A), barrier tissues (B), and metabolically active tissues (C). Percentages indicate the frequencies of ST2⁺ and ST2⁻ populations within the Treg cell population (left plots), and the frequency of CXCR3⁺ cells within ST2⁺ (pink) and ST2⁻ (light blue) Treg cell subsets (right plots). (D) Summary plot of the analyzed steady-state tissues showing the frequency of ST2⁺ Treg cells and the proportion of CXCR3⁺ cells within the ST2⁺ Treg cell compartment.

**Supplemental Figure S5.**
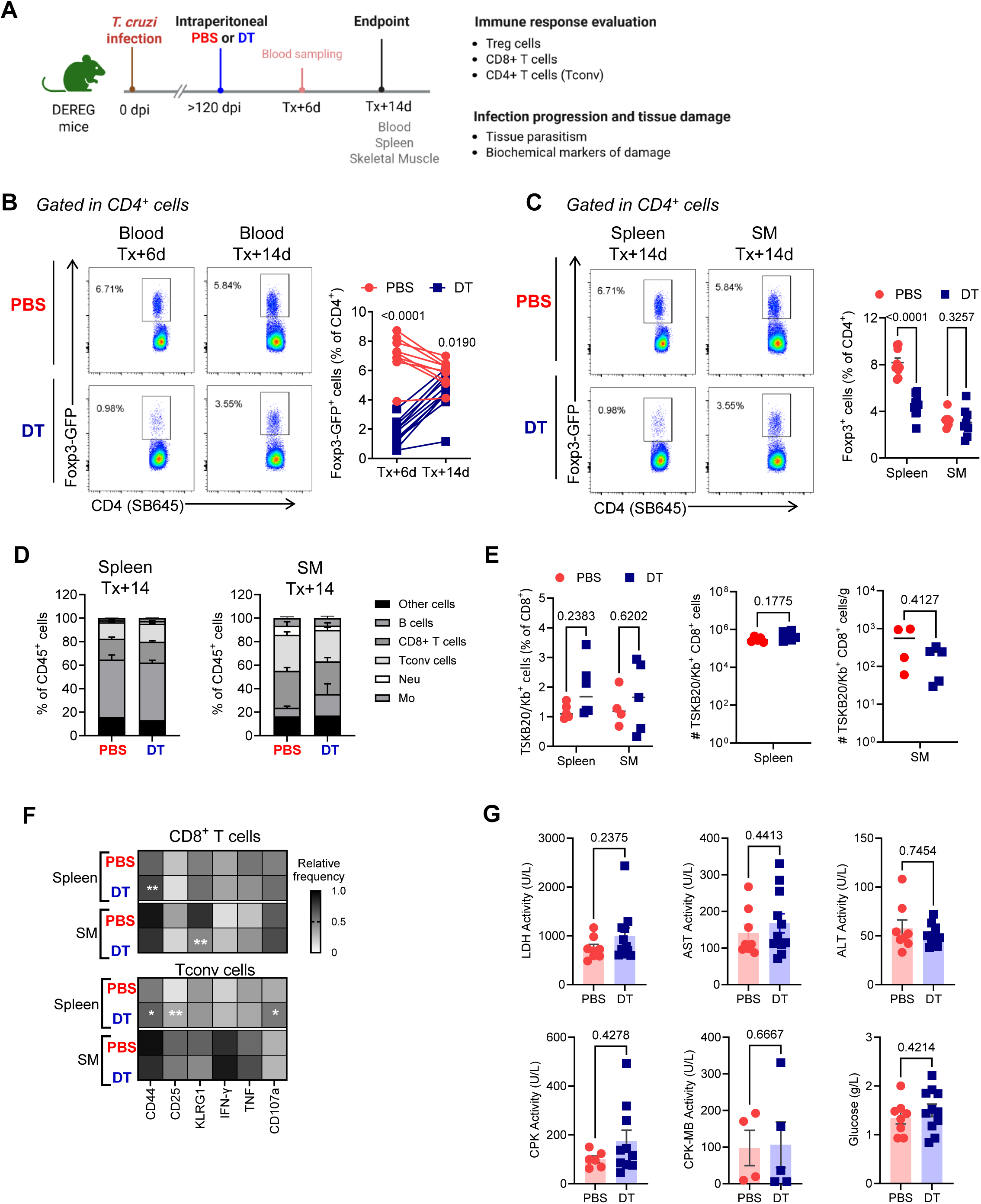
Transient systemic Treg cell depletion has limited impact on immune responses and tissue damage during chronic infection. Chronically infected mice received a single intraperitoneal administration of diphtheria toxin (DT) or PBS at ≥120 days post infection (dpi). Blood was collected at 6 and 14 days after treatment (Tx+6d and Tx+14d), whereas spleen and skeletal muscle (SM) were collected at Tx+14d for analyses. (A) Experimental scheme. (B) Representative flow cytometry plots and paired analysis of circulating Foxp3⁺ Treg cell frequencies. (C) Representative flow cytometry plots and summary quantification showing Foxp3⁺ Treg cells in spleen and SM. (D) Relative composition of major lymphoid and myeloid cell populations in spleen and SM. (E) Frequency and absolute numbers of parasite-specific TSKB20/Kb^+^ cells within CD8^+^ T cells in spleen and SM. (F) Heat map summarizing changes in activation and effector markers expression within Foxp3^-^CD4⁺ (T conv) and CD8⁺ T cell populations. (G) Plasma levels of biochemical markers of tissue damage. Across panels, each symbol represents one mouse. Bars indicate mean ± SD where applicable. Statistical analyses were performed as described in the Methods.

**Supplemental Figure S6.**
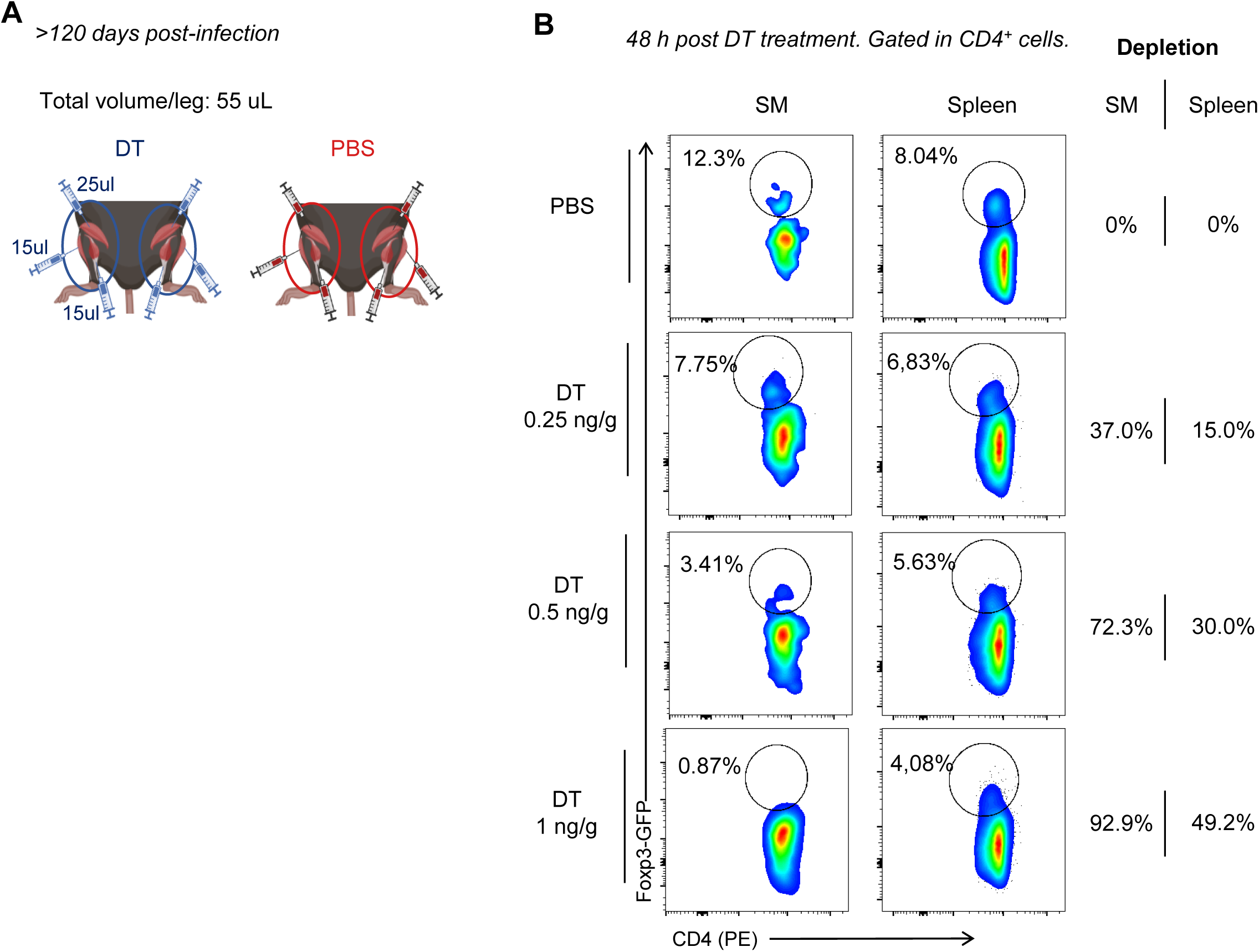
Optimization of a protocol for local Treg cell depletion in skeletal muscle. (A) Schematic representation of intramuscular administration of PBS or diphtheria toxin (DT) into posterior limb muscles (quadriceps, gastrocnemius, and tibialis anterior) of chronically infected DEREG mice. (B) Representative flow cytometry plots showing gating of Foxp3⁺ Treg cells in skeletal muscle and spleen 48 h after intramuscular administration of PBS or DT at the indicated doses. Numbers indicate the percentage of Treg cell depletion relative to PBS-injected controls.

**Supplemental Figure S7.**
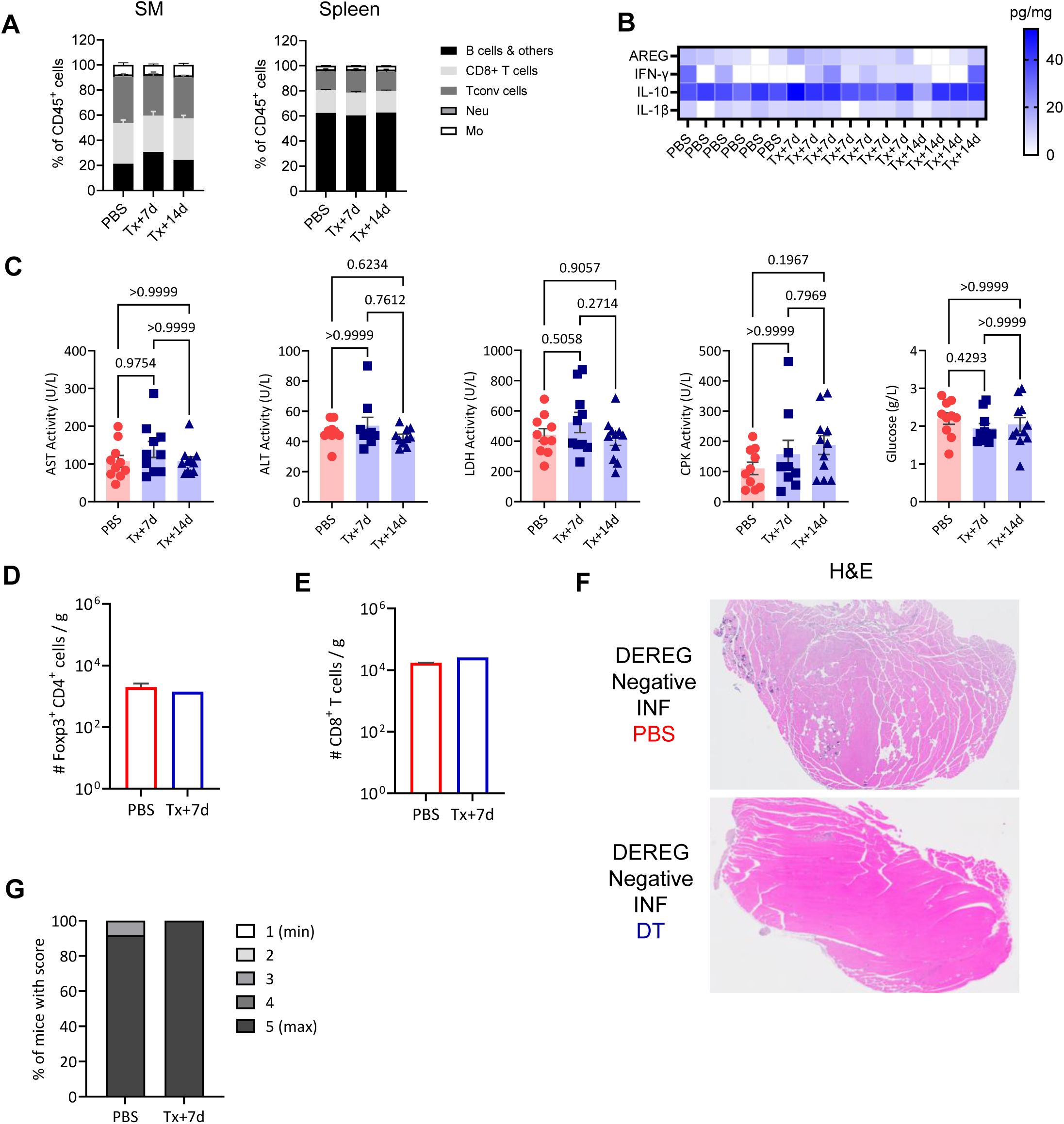
Local Treg cell depletion does not alter major immune populations or cytokine levels and DT administration has no detectable effects in DEREG-negative littermates. (A-C) Chronically infected DEREG mice received intramuscular injections of PBS or diphtheria toxin (DT) into posterior limb muscles (quadriceps, gastrocnemius, and tibialis anterior) at ≥120 dpi. DT-treated mice were analyzed at Tx+7d and Tx+14d. PBS-treated mice from both time points were pooled and analyzed as a single control group. (A) Relative composition of major lymphoid and myeloid cell populations skeletal muscle (SM) and spleen. (B-C) Cytokines concentration (B) and biochemical markers of tissue damage (C) quantified in plasma. (D–G) Control experiments in chronically INF DEREG-negative littermates receiving local intramuscular PBS or DT at Tx+7d. (D) Absolute numbers of Foxp3⁺ Treg cells in SM. (E) Absolute numbers of total CD8⁺ T cells in SM. (F) Representative hematoxylin and eosin–stained (H&E) sections of SM. (G) Skeletal muscle performance assessed by Kondziela’s inverted screen test. Across panels, each symbol represents one mouse. Bars indicate mean ± SD where applicable. Statistical analyses were performed as described in the Methods.

**Supplemental Figure S8.**
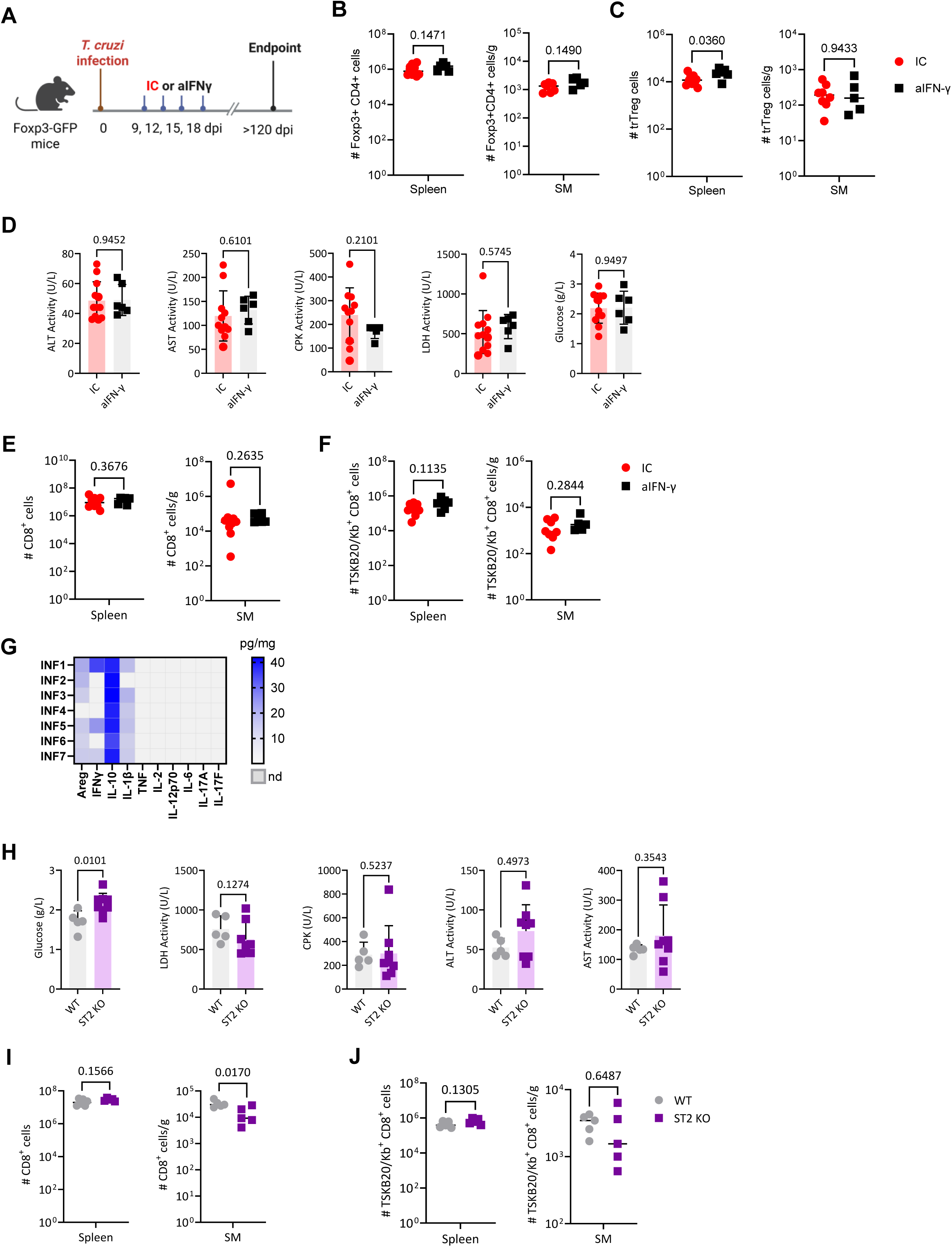
Effects of acute IFN-γ blockade and ST2 deficiency during chronic T. cruzi infection. (A-F) *T. cruzi* infected Foxp3-GFP mice were treated with anti–IFN-γ or isotype control during the acute phase and analyzed at ≥120 dpi. (A) Experimental scheme. (B-C) Absolute numbers of total Foxp3⁺ Treg cells (B) and tissue-repair Treg (trTreg) cells (C) in spleen and SM. (D) Plasma biochemical markers of tissue damage. (E-F) Absolute numbers of total (E) and parasite-specific TSKB20/Kb⁺ (F) CD8⁺ T cells in spleen and skeletal muscle (SM). (G) Heat map showing relative levels of cytokines detected in SM from INF mice. Each row represents one mouse. (H-J) Wild type (WT) and ST2-deficient mice (ST2 KO) were infected with *T. cruzi* and evaluated at ≥120 dpi. (H) Plasma levels of biochemical markers of tissue damage. (I-J) Absolute numbers of total (I) and parasite-specific TSKB20/Kb⁺ (J) CD8⁺ T cells in spleen and SM. Across panels, each symbol represents one mouse. Bars indicate mean ± SD where applicable. Statistical analyses were performed as described in the Methods.

## Supplemental Methods

**Supplemental Table 1.** Antibodies and reagents.

| Flow cytometry reagents |  |  |  |  |  |
| --- | --- | --- | --- | --- | --- |
| Marker | Fluorophore | Clone | Vendor | Catalog # | Dilution |
| Areg | Biotinilayed | Polyclonal | R&D Systems | BAF989 | 1/50 |
| B220 | PE-Cy7 | RA3-6B2 | Invitrogen | 25-0452-82 | 1/300 |
| B220 | Brilliant Violet 711 | RA3-6B2 | BD | 563892 | 1/300 |
| B220 | APC-Cy7 | RA3-6B2 | Biolegend | 103224 | 1/200 |
| CD107a | PE | 1D4B | Biolegend | 121612 | 1/200 |
| CD11b | PerCpCy5.5 | M1/70 | Biolegend | 101228 | 1/100 |
| CD11b | Super Bright 645 | M1/70 | Invitrogen | 64-0112-80 | 1/200 |
| CD11b | eFluor 660 | M1/70 | eBioscience | 50-0112-82 | 1/100 |
| CD25 | PE-Cy7 | PC61.5 | eBioscience | 25-0251-82 | 1/50 |
| CD3e | APC | 145-2C11 | eBioscience | 17-0031-82 | 1/100 |
| CD4 | Super Bright 645 | GK1.5 | Invitrogen | 64-0041-82 | 1/200 |
| CD4 | PE | GK1.5 | eBioscience | 12-0041-83 | 1/400 |
| CD4 | APC-eFluor 780 | GK1.5 | invitrogen | 47-0041-82 | 1/100 |
| CD4 | PE-Cy7 | GK1.5 | eBioscience | 25-0041-82 | 1/200 |
| CD4 | APC-eFluor 780 | GK1.5 | invitrogen | 47-0041-82 | 1/200 |
| CD44 | PE-Cy5 | IM7 | eBioscience | 15-0441-81 | 1/300 |
| CD45 | Alexa Fluor 700 | 30-F11 | Thermofisher | 56-0451-82 | 1/100 |
| CD45 | PE-Cy7 | 30-F11 | eBioscience | 25-0451-82 | 1/400 |
| CD45 | APC-Cy7 | 30-F11 | BD | 557659 | 1/400 |
| CD62L | Super Bright 600 | mel-14 | eBioscience | 63-0621-82 | 1/200 |
| CD62L | Super Bright 600 | mel-14 | eBioscience | 63-0621-82 | 1/200 |
| CD8a | PE-Cy5.5 | 53-6.7 | eBioscience | 35-0081-82 | 1/300 |
| CTLA4 | Brillant Violet 605 | UC10-4B9 | Biologend | 106323 | 1/100 |
| CXCR3 | Brillant Violet 510 | CXCR3-173 | Biologend | 126528 | 1/100 |
| GzmB | APC/Fire 750 | QA16A02 | Biologend | 372210 | 1/75 |
| H-2Kb ANYKFTLV (TSKB20) | Brillant Violet 421 |  | NIH Tetramer Core Facility | 52270 H-2K (b) | 1/300 |
| IFN-γ | Brillant Violet 711 | XMG1.2 | Biologend | 505836 | 1/100 |
| IFN-γ | Brillant Violet 711 | XMG1.2 | Biologend | 505836 | 1/100 |
| KLRG-1 | PE-eFluor 610 | 2F1 | eBioscience | 61-5893-82 | 1/300 |
| LIVE/DEAD Fiable Near-IR Dead Cell Stain Kit |  |  | Invitrogen | L10119 | 1/1000 |
| LIVE/DEAD™ Fixable Aqua Dead Cell Stain Kit for 405 nm excitation |  |  | Invitrogen | L34966 | 1/500 |
| Ly6G | PE-Cy7 | 1A8 | Biologend | 127618 | 1/200 |
| Ly6G/Ly6C | Super Bright 702 | RB6-8C5 | Invitrogen | 67-5931-82 | 1/150 |
| PD-1 | PE | J43 | eBiosciences | 12-9985-81 | 1/200 |
| ST2 | Brillant Violet 785 | DIH9 | Biologend | 145321 | 1/25 |
| ST2 | PE | DIH9 | Biologend | 145304 | 1/25 |
| Streptavidin | Qdot605 |  | Invitrogen | Q10101MP | 1/200 |
| T-bet | eFluor 660 | eBio4B10 | eBioscience | 50-5825-82 | 1/75 |
| TNF | eFluor 450 | MP6-XT22 | Invitrogen | 48-7321-82 | 1/100 |
| TNF | PerCp-Cy5.5 | MP6-XT22 | Biologend | 506322 | 1/100 |

| Other reagents |  |  |  |  |
| --- | --- | --- | --- | --- |
| Product | Assay | Vendor | Catalog # | Observaciones |
| TRIzol Reagent | DNA extraction | Invitrogen | 15596018 |  |
| Custom TaqMan Gene Expression Assay targeting *T. cruzi* satellite DNA | qPCR | Applied Biosystems | 4332078 |  |
| TaqMan Rodent GAPDH Control Reagents | Endogenous control qPCR | Applied Biosystems | 4308313 |  |
| TaqMan Universal Master Mix II, no UNG | qPCR | Applied Biosystems | 4440040 |  |
| LEGENDplex™ Mouse Th Cytokine Panel (12-plex) | Multiplex cytokine assay | Biolegend | 741043 | IFN-γ, IL-5, TNF, IL-2, IL-6, IL-4, IL-10, IL-9, IL-17A, IL17F, IL-22, IL-13 |
| LEGENDplex™ Mouse Cytokine Panel 2 (13-plex) | Multiplex cytokine assay | Biolegend | 741335 | IL-12p70, IL-1α, IL-1β, IL-3, IL-12p40, IL-23, IL-7, IL-11, IL-27, IFN-β, GM-CSF, TSLP |
| Mouse IL-33 ELISA Kit | ELISA | Invitrogen | BMS6025TEN | Quantification of IL-33 protein |
| Dyphtheria Toxin | Treg depletion | Calbiochem | 322326 |  |
| Mouse Amphiregulin DuoSet ELISA | ELISA | R&D Systems | DY989 |  |
| *InVivo*MAb anti-mouse IFN-γ | Blocking | Bio X Cell | #BE0055 | clone XMG1.2 |
| *InVivo*MAb rat IgG1 isotype control, anti-horseradish peroxidase | IFN-γ neutralization control | Bio X Cell | #BE0088 | Clone HRPN |

## Notes

### Competing Interest Statement

The authors have declared no competing interest.

